# Quantifying isolation-by-resistance and connectivity in dendritic ecological networks

**DOI:** 10.1101/2021.03.25.437078

**Authors:** Tyler K. Chafin, Steven M. Mussmann, Marlis R. Douglas, Michael E. Douglas

## Abstract

1. A central theme in landscape ecology is the translation of individual movements within a population by deconstructing/interpreting the components of its topographical environment. Most such endeavors rely heavily on the concept of ’landscape resistance’ – a composite of an arbitrary number of features/covariates that, when identified/compiled, yield a ‘surface’ inversely related to net movement. However, the statistical methodologies underlying this compilation have limited applicability when applied to dendritic ecological networks (DENs), including riverscapes.
2. Herein we provide an analytical framework (ResistNet) that more appropriately annotates DEN segments by first aligning individual genetic distances with environmental covariates within a graph structure, then employing a genetic algorithm to optimise a composite model.
3. We evaluated the efficacy of our method by first testing it *in silico* across an array of sampling designs, spatial trajectories, and levels of complexity, then applying it in an empirical case study involving 13,218 ddRAD loci from N=762 Speckled Dace (Leuciscidae: *Rhinichthys osculus*), sampled across N=78 Colorado River localities. By doing so, we underscored the utility of ResistNet within a large-scale conservation study, as well as identified prerequisites for its appropriate application.
4. Our contemporary framework not only allows an interpretation of meta-population/meta-community structure across DENs, but also highlights several innovative applications. These are: (a) Expanding an ongoing study design, and thus its hypotheses, into yet unsampled temporal and/or spatial arenas, and; (b) Promoting multi-species management through comparative analyses that extend across species and/or drainages.

## 1. INTRODUCTION

The physical structure of watersheds not only defines form and function in freshwater fishes (Douglas & Matthews, 1992; Jackson et al., 2001), but also their intraspecific evolutionary trajectories and population dynamics (Hopken et al., 2013). As a result, patterns of diversification differ markedly from those found in terrestrial land- and seascapes (Manel et al., 2020; Martinez et al., 2018). The unique constraints of riverine networks serve not only to define dispersal pathways but also to dictate genetic relatedness within/ among aquatic biota (Meffe & Vrijenhoek, 1988). Network ‘connectivity’ (=potential for movement), and ‘complexity’ (=branching rates/ nonlinear topology) control numerous population processes, such as: Genetic divergence (Hopken et al., 2013), colonization probabilities (Falke et al., 2012; Labonne et al., 2008), and metapopulation components (Campbell Grant, 2011; Fagan, 2002).

In general, complex dendritic networks may promote intraspecific genetic diversities (Thomaz et al., 2016), and divergences among distal populations (Chiu, Li, et al., 2020). Yet, the unidirectional flows in riverscapes differ from other such dendritic ecological networks (DENs), and thus biases dispersal (i.e., up *versus* downstream). This in turn promotes asymmetric gene flow and source-sink metapopulation structure (Campbell Grant et al., 2007). Allelic diversity increases in a downstream direction (Blanchet et al., 2020; Paz-Vinas et al., 2015; Paz-Vinas & Blanchet, 2015), whereas unique genetic variation is most often retained within peripheral headwater populations (Chiu, Nukazawa, et al., 2020; Morrissey & De Kerckhove, 2009).

Conversely, effective population size (Altermatt & Fronhofer, 2018) and genetic variability (Thomaz et al., 2016) seemingly decrease within unimpeded linear systems which, in turn, impacts both resilience and stability of lineages (Mari et al., 2014; Terui et al., 2019), as well as proliferation capacity (Harvey et al., 2019). However, numerous considerations can alter these expectations, such as: Dispersal bias (Chiu, Li, et al., 2020), nonequilibrium processes (Pilger et al., 2017), geomorphic/ climatic regimes (Douglas et al., 1999; Osborne et al., 2014), and abiotic species-level interactions (Paz-Vinas et al., 2015; Rodríguez-González et al., 2019).

It is thus not surprising that evaluation of stream distances alone rarely captures intraspecific genetic patterns. In a comparison of 12 salmonid species, isolation-by-distance (IBD) varied from *r*^2^=0.06–0.87 (Bradbury & Bentzen, 2007), and from 0.0001–0.44 among three darter species (Camak & Piller, 2018). A larger-scale study involving 31 co-distributed stream fishes across 7 families identified a significant relationship between among-site *F*_st_ and log-transformed river network distances (Mantel *r*=0.11–0.88), with stream hierarchy explaining the greatest proportion of genetic variance (Zbinden et al., 2022a). Although such deviations are partly driven by stochastic or nonequilibrium processes (Hutchison & Templeton, 1999), their residuals should also reflect spatial features that impede individual movements (Keis et al., 2013; Tang et al., 2020). Thus, divergence can be considered as a general function of the movement-cost along stream segments, i.e., ‘isolation-by-resistance’(IBR)(B. H. McRae, 2006; B. H. McRae & Beier, 2007).

The IBR model applied within dendritic networks emphasizes specific attributes of individual edges or stream segments, with resistance amplified by an array of localized factors (Fullerton et al., 2010; Hughes et al., 2009): Rate of elevational change (Lowe et al., 2006), flow regime (Rolls et al., 2016), channel width/ catchment area (Ma et al., 2020), water composition (Fourtune et al., 2016), and seasonal hydroclimate (Brauer et al., 2016, 2018). In addition, organisms that never (or rarely) occupy upstream or headwater conditions can still be directly affected by those habitats (Frissell et al., 1986; Fullerton et al., 2010; Nadeau & Rains, 2007).

Most linear streams are inherently predisposed to fragmentation (Brauer & Beheregaray, 2020; Fagan et al., 2002; Fagan, 2002), a situation particularly exacerbated, given that relatively few remain free-flowing (Grill et al., 2019). Water storage also impacts habitat suitability (Chafin et al., 2019; Fraser et al., 2019), with populations shifting trajectories accordingly (Dibble et al., 2021). Dams and diversions extirpate populations and promote homogenization by disrupting homing behaviors (Baggio et al., 2018), with species of limited mobility particularly impacted (Coleman et al., 2018). Again, vulnerability and the specific nature of impacts vary by context and species (Prunier et al., 2018).

### 1.1 Landscape genetic methods applied to riverscapes

The mechanics of landscape genetics/genomics often translate poorly to dendritic networks (Davis et al., 2018; Grummer et al., 2019). At issue is the riverscape structure upon which genetic relationships can be mapped. Frequent *ad hoc* approaches often lack an explicit spatial framework, or rely upon spatial methodology is landscape-based (Cooke et al., 2014; Galbraith et al., 2015; Kanno et al., 2011). Often, these methods do so by contrasting a genetic distance matrix against pairwise differences in local conditions, often using the Mantel test and derivatives (Douglas & Endler, 1982; Smouse et al., 1986), thus neglecting the space through which individuals must travel between points, restricting their riverscape-applicability (Davis et al., 2018).

On the other hand, path-based methods (as herein) consider the cumulative costs of movement, with resistance being represented as a composite of contributing environmental factors (Zeller et al., 2012) that allow for habitat-related metrics to be derived, such as: Least-cost paths, net flux between points, and movement(B. McRae et al., 2016). Operationally, it involves summarizing spatial variables as a raster image of *n* x *m* cells (analogous to pixels), with assigned values representing ‘resistance’ (Davis et al., 2018).

We propose a graph structure to translate this concept to riverscapes, with edges annotated with environmental attributes connecting the nodes (i.e., populations or junctions) (Eros et al., 2011; Peterson et al., 2013). Few approaches have considered resistance within a spatially explicit, graph-theoretic framework, and even fewer have incorporated multiple covariates within an hypothesis-testing framework [although see (White et al., 2020)]. Here, an heuristic approach that explores complex parameter space is required, particularly given the sensitivity of resistance to parameterization, and the difficulties with the *a priori* weighting of specific variables within a larger pool of candidates (Spear et al., 2010). These approaches have previously been developed for landscapes, using algorithms for model exploration and selection (Peterman, 2018), and proven successful in both empirical and *in silico* scenarios (Kimmig et al., 2020; Peterman et al., 2019; Winiarski et al., 2020).

Here, we employ a similar resistance perspective to quantify riverscape genetics/ genomics and do so by exploiting large-scale, contemporary efforts that quantify/ classify stream variables (Grill et al., 2019; Linke et al., 2019; McManamay et al., 2018; Troia & McManamay, 2020). We utilised our package (ResistNet) to evaluate a case study involving Speckled Dace (Leuciscidae, *Rhinichthys osculus*), the most widely distributed freshwater fish endemic within the most endangered and regulated river system of North America, the Colorado River (Grill et al., 2019; Minckley & Deacon, 1968; Nilsson et al., 2005). To accomplish this, we analyzed a large ddRAD dataset to test the following hypotheses: 1) Populations are spatially structured within the topology of the dendritic network; 2) Genetic differentiation is modulated by environmental attributes, representing edges within the river network; and 3) Spatio-genetic patterns cannot be explained by distance alone, but rather by a model of IBR.

## 2. METHODS

ResistNet facilitates high-throughput analyses by automating the association between environmental correlates and genetic differentiation in dendritic networks (Figure 1). Source code and documentation is available at: github.com/tkchafin/resistnet.

**Figure 1:**
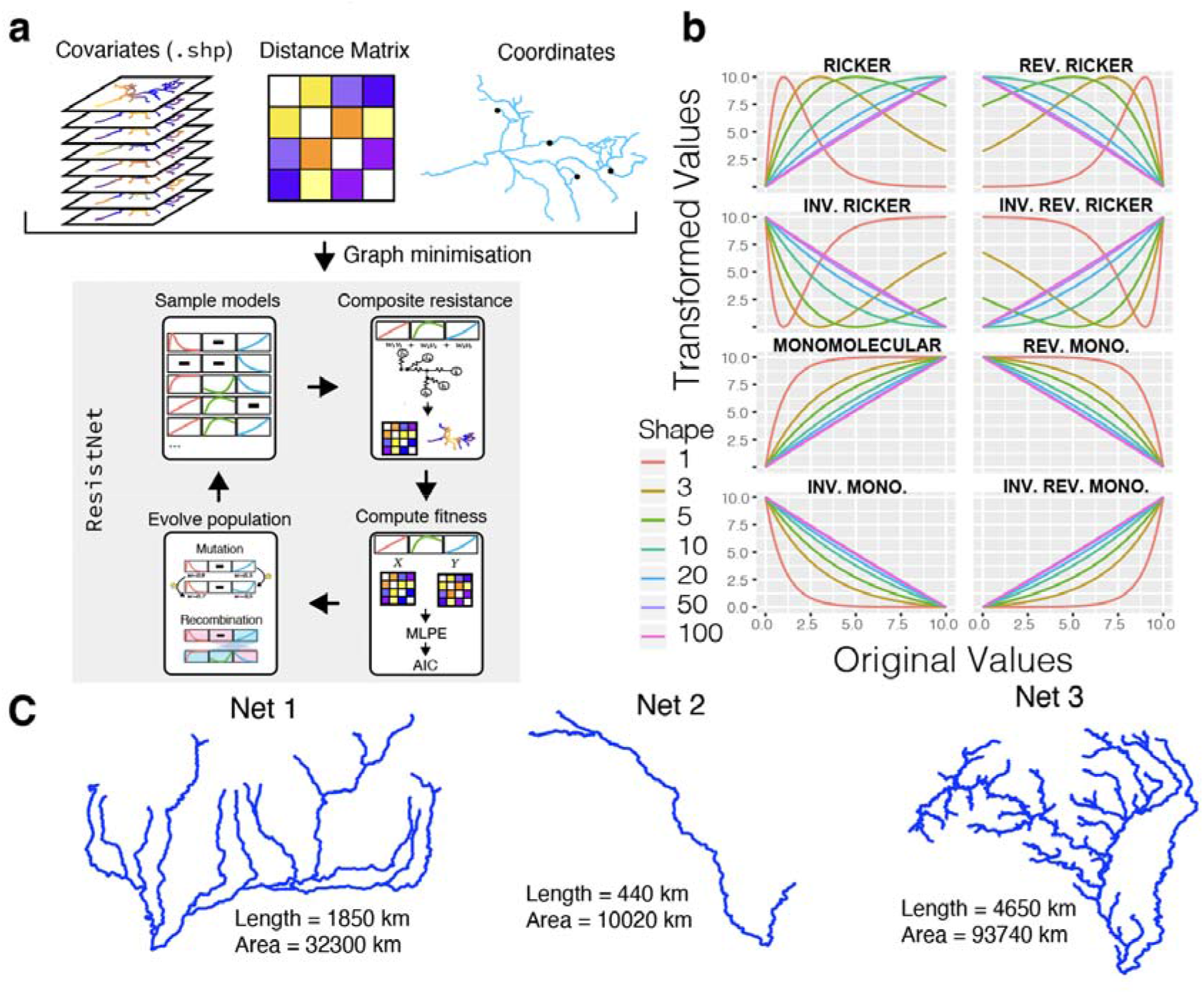
Overview of ResistNet and its validation environments. (a) Inputs consist of a spatial network with *n* covariates encoded as reach-level attributes (as shapefile, *.shp*), and pairwise genetic distances for georeferences samples. These are used to construct a minimal graph representation as the primary data structure of ResistNet. A population of models randomly initialises the genetic algorithm, with fitness computed using a Maximum-Likelihood Population Effects (MLPE) model applied to the composite, transformed ‘effective’ resistance values. Optimisation proceeds iteratively following random sampling (mutation) of parameter values, swapping between models (recombination), and ‘selection’ to generate offspring. (b) Variable transformation occurs given eight functions, or a subset thereof (per user configuration). (c) Three input networks of varying topological complexity and spatial extent served as templates from which to simulate distance matrices to test ResistNet (Net1-3).

### 2.1 Composite resistance models optimized within networks

ResistNet combines a genetic algorithm with multiple environmental variables as a mechanism to parameterize resistance (Figure 1), similar to that found in ResistanceGA (Peterman, 2018), but with several important differences. Genetic algorithms are useful for model optimisation and selection of features in that they are less impacted by local optima when compared to other hill-climbing techniques (Prügel-Bennett, 2004). They are employed to simultaneously evaluate a *population* of models by using log-likelihood or AIC to gauge *fitness*. Here, *individuals* (as randomly-sampled model configurations) consist of *chromosomes*, with *genes* (as parameter choices) represented by 1) Variable inclusion/exclusion; 2) Variable transformations (using functions available in ResistanceGA; Supplementary Figure S1); 3) Shape parameters for transformation; or 4) Variable weights (with user-defined ranges).

For an optimisation problem involving *n*-environmental parameters, an individual chromosome has length 4*n*. Fitness values (e.g., AIC) then determine the survival of individual models and their contribution to successive generations, with stochasticity represented by *mutation* (random changes to parameter values) and *recombination* (parameter choices by multiple individuals combined as offspring). Both of these occur following user-defined probabilities. Optimisation continues until a threshold number of generations have passed without an increase in population fitness.

The workflow used by ResistNet (Figure 1) is summarized as follows:

1. Network structure and environmental data are imported as shapefiles. Environmental data for contiguous stream reaches (=edges) are summarised and rescaled from 0–10, (per Peterman, 2018).
2. The initial population has a user-defined size and randomly seeded, which initialises the genetic algorithm (as implemented using DEAP; De Rainville et al., 2012). Individuals (as chromosomes) contain transformation, shape, and weight parameters to ensure an optimal scale per a potential non-linear relationship between gene flow and each variable. This process also allows the relationship in a composite resistance model to vary according to relative influence and directionality (e.g., positive/ negative).
3. Cumulative effective resistance is then computed pairwise between nodes as a weighted sum of (transformed/ scaled) individual covariates (McRae et al., 2008; Thiele et al., 2018). By default, each covariate is individually aggregated pairwise between samples as the sum of values along the Dijkstra least-distance path, although custom behaviours can be user-specified. This results in a single composite pairwise resistance matrix for each model.
4. Individual fitness is calculated via a mixed effects maximum likelihood population effects (MLPE) model (Clarke et al., 2002; Van Strien et al., 2012). This allows the pairwise genetic and resistance matrices to be compared and fitted using the MLPE function from ResistanceGA. Pairwise distances are inherently non-independent, a condition MLPE models were specifically formulated to accommodate, and are thus effective for landscape genetic model selection (Peterman & Pope, 2021; Shirk et al., 2018). Fitness for the genetic algorithm is assessed as log-likelihood, AIC, or marginal *R*^2^, but may also be computed using ΔAIC_null_ (i.e., the change in AIC values as compared to a random effects null model).
5. Individuals are selected for successive generations via ‘tournament selection,’ a deterministic method whereby random sets of individuals (user-specified size *n*) are compared (=tournaments), with ‘winners’ being retained.
6. Mutation (=changing parameter values) and recombination (=exchanging parameter combinations among models) occur at generational intervals, with the respective probabilities of each being user-specified.
7. Steps 2-6 are repeated for a user-specified number of generations, or until user-specified generations fail to increase the maximum fitness beyond a given threshold.
8. After stopping criteria are met, Akaike weights are computed for the highest fitness models in the population, with these being retained to meet a cumulative user-specified Akaike weight threshold (default value=0.95). Model-averaged resistance and multi-model relative variable importance values are subsequently computed using the chosen model set (below).

Multi-model inference has several benefits: It avoids the inherent subjectivity of model selection, in that estimates from every candidate model are integrated proportional to their approximation of the observed genetic distances [using the Akaike weight (*w*_i_ for *i* models)], while also more concisely incorporating both uncertainty and information from closely ranked models. Additionally, it also allows the quantification of relative variable importance (RVI) across models composed of varying numbers of explanatory variables, and does so by deriving the sum of Akaike weights (SW) as a robust measure of RVI in a multi-model inference framework (Giam & Olden, 2016).

### 2.2 Simulation-based validation

To validate the capacity of ResistNet to accurately recover a known model, we employed a simulation scheme involving three river networks of varying complexity (near-linear, moderately branched, and highly branched; Figure 1) and four scenarios involving different environmental covariates (Table 1), to include distance-only models, and those varying in weights. Three sampling levels (10, 30, and 50) were evaluated, with 20 replicates per level. Simulations were conducted using *simResistNet.py*, an auxiliary software that simplifies validation with a specific riverscape and its combination of environmental variables. Input consists of shapefiles for the three template networks, and simulation parameters (as tab-delimited text) specifying covariates for inclusion, their weights, and any transformations that must be applied. These are employed to compute pairwise resistance distances between sampled points, which are then normalised between 01.

**Table 1:** RiverATLAS variables used as covariates for modeling effective resistance networks in ResistNet. Inclusion as potential covariates simulation-based validation and empirical case study in *Rhinichthys osculus* is indicated as “*”. Weights (*w*) are provided for each of the simulated scenarios (Sim1-4).

| Var. | Description | Empirical | Simulation | wSim1 | wSim2 | wSim3 | wSim4 |
| --- | --- | --- | --- | --- | --- | --- | --- |
| AET | Actual evapotranspiration; equivalent to ‘soil water stress coefficient’ | * | - | - | - | - | - |
| ARI | Aridity index | * | - | - | - | - | - |
| CRP | Cropland extent as percentage of reach catchment | * | - | - | - | - | - |
| DOR | Degree/ index of anthropogenic flow regulation | * | * | - | - | 0.5 | 0.5 |
| FOR | Forest landcover percentage in reach catchment | * | - | - | - | - | - |
| HFT | Standardized human footprint index | * | - | - | - | - | - |
| KAR | Percentage of catchment exposed karst terrain | * | - | - | - | - | - |
| LEN | Reach length in kilometers | * | - | 1.0 | - | - | - |
| LKA | Limnicity (percent lake area in reach catchment) | * | - | - | - | - | - |
| PAC | Percentage of reach catchment in protected areas | * | - | - | - | - | - |
| POP | Scaled human population size in 2010 | * | - | - | - | - | - |
| PRE | Precipitation, as annual average in reach catchment | * | * | - | - | - | - |
| PST | Pasture extent as percentage of reach catchment | * | - | - | - | - | - |
| RDD | Road crossing density | * | * | - | - | - | 0.5 |
| RIV | River volume along reach segment | * | - | - | - | - | - |
| REV | Reservoir volume in watershed upstream of reach | * | - | - | - | - | - |
| RUN | Annual average land surface discharge and runoff | * | - | - | - | - | - |
| SGR | Stream gradient; ratio of elevation differential in reach and length | * | - | - | 1.0 | 1.0 | 1.0 |
| SLP | Average slope degree in catchment | * | - | - | - | - | - |
| SOC | Soil organic carbon content | * | - | - | - | - | - |
| TMP | Mean annual temperature | * | * | - | 1.0 | 1.0 | 1.0 |
| URB | Urban extent | - | * | - | - | - | - |

ResistNet was run at 600 generations maximum on each replicate, with an ending condition of 20 generations of continuous failure to increase maximum observed fitness, a fixed population size of N=600, a variable weight floor of N=0.2, a maximum transformation shape of n=0,5, and mutation/ recombination probabilities of 0.8. To facilitate replication, the evaluation procedure is implemented as a Snakemake workflow (Mölder et al., 2021), enabling users to apply it to be applied by users to their own input networks and environmental models.

We examined model-averaged weights (MAW) and multi-model relative variable importance values (RVI) to gauge performance metrics against input conditions. These were assessed in three ways: (1) Summarizing among-replicate variation using median RVI and mean MAW; (2) Re-computing model averages as an Akaike-weighted ensemble model, including all sampled models among replicates; and (3) Re-computing model averages using only the best AIC model from each replicate run. For ensemble models, only models below a cumulative Akaike threshold of 0.95 were retained.

### 2.3 Rhinichthys osculus *as a case study*

We focused on the small-bodied *Rhinichthys osculus*, broadly endemic to western North America within all major drainage basins west of the Rocky Mountains (Hubbs & Miller, 1948; Minckley et al., 1986; Oakey et al., 2004; Smith & Dowling, 2008). As such, it is remarkably adaptive, with a range extending from high-altitude/ cold-water Pacific Northwest streams (Pfrender et al., 2004; Wiesenfeld et al., 2018), to endorheic basins vulnerable to stochastic and anthropogenic influences in the Mojave/ Sonoran deserts (Mussmann, Douglas, Oakey, et al., 2020). Click or tap here to enter text.Its extinction vulnerability is underscored by the contemporaneous elimination of two lineages within the 20^th^ century, and four additional subspecies listed under the United States Endangered Species Act of 1973, as amended (16 U.S.C. 1531-1544, 87 Stat. 884). Our case study explores the utility of ResistNet in categorizing extant conservation and management challenged for this species, with a focus on distributions localised within the Colorado River Basin.

A matrix of unlinked single nucleotide polymorphisms (SNPs) was derived from existing datasets (Mussmann, 2018). Briefly, genomic DNA was extracted from fin tissue non-lethally sampled from *N*=762 individuals distributed throughout 78 Colorado River Basin localities (mean=9.8/site) (Figure 2; Supplementary Table S1; Mussmann, 2018). Library preparation followed the ddRAD protocol (Peterson et al. 2012), using *Pst*I and *Msp*I enzymes (New England Biolabs) with 144 samples pooled per 1x100bp lane for sequencing on the Illumina HiSeq 2000 (University of Wisconsin Biotechnology Center), or HiSeq 4000 (University of Oregon Genomics & Cell Characterization Core Facility).

**Figure 2:**
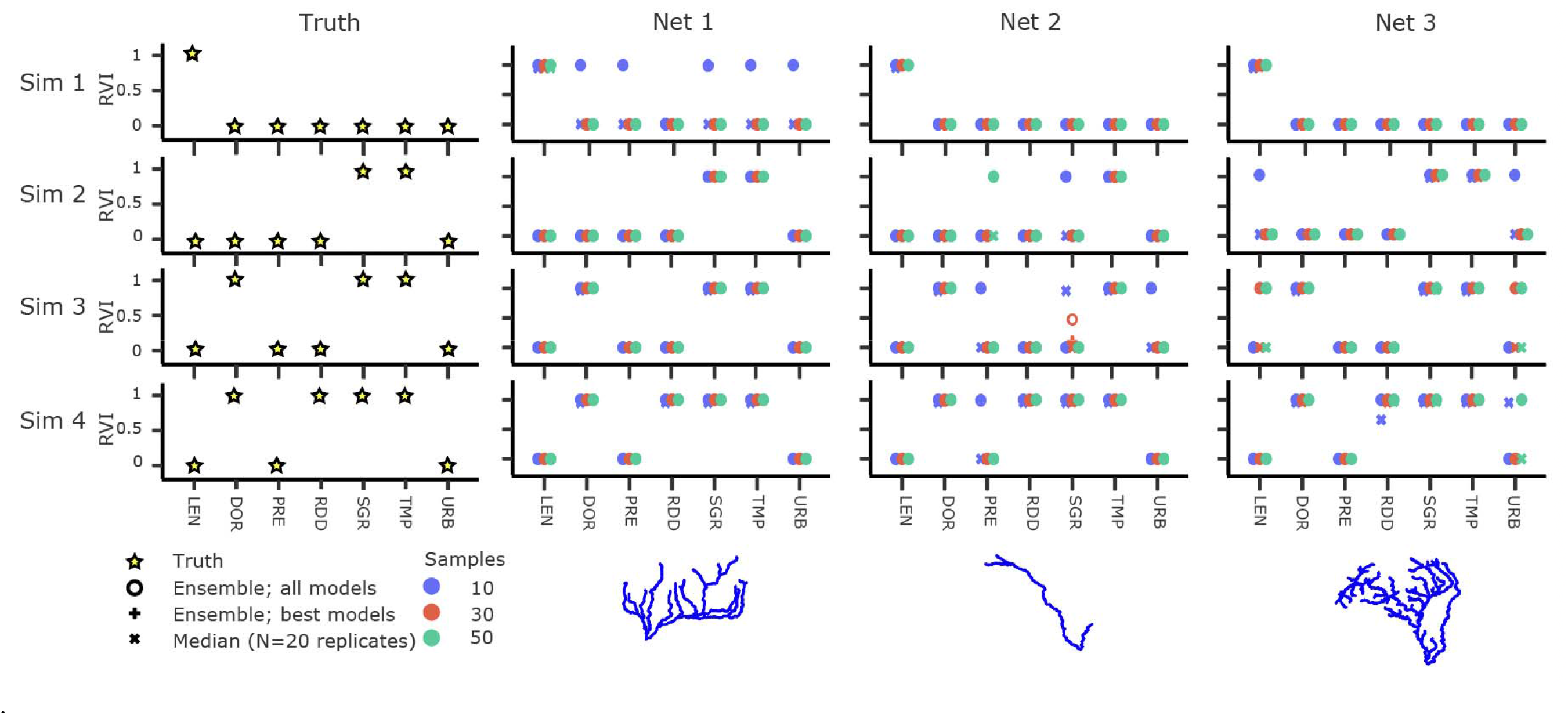
Summary of recovered relative variable importance (RVI) from simulated scenarios. RVIs were derived from four pre-determined ‘known’ scenarios (Sim1-4), tested within three distinct template networks (Net1-3). Results are reported as: (1) Ensemble (all): a model-averaged RVI computed from all models chosen across 20 independent replicates; (2) Ensemble (best): A model-averaged RVI calculated exclusively from the ’optimum’ models in each replicate, determined by the Akaike Information Criterion (AIC); and (3) Median: The median RVI amongst all replicates. Each scenario was modeled using three different sample sizes: 10, 30, and 50, represented by blue, red, and green points, respectively.

Reads were demultiplexed and assembled via *de novo* clustering in Stacks v.2.41 (Catchen et al., 2013). Samples were grouped hierarchically into *N*=5 subregions: Upper Colorado River Basin (*N*=221); Virgin River (*N*=223); Grand Canyon and tributaries (*N*=149); Lower Colorado River Basin (*N*=105); and Little Colorado River (*N*=64). Trial runs using STACKS yielded optimal parameters (Paris et al., 2017) for minimum number of raw reads needed per stack (m=4), as well as number of mismatches between stacks for identification of putative loci (M=2), and catalog formation (n=2). Loci appearing in <50% of individuals per subregion were removed, as were loci with an observed heterozygosity >0.6, and a minor allele frequency (MAF) <0.01. These measures effectively removed putative paralogs and minimally informative loci (Eaton, 2014; McKinney et al., 2017).

#### 2.3.1 Population structure

To provide context for riverscape resistance and genetic differentiation, we first gauged population structure using an aspatial clustering method (ADMIXTURE v1.3.0; Alexander et al., 2009). Data were filtered to one SNP/ locus, with samples partitioned by region. In doing so, we first evaluated models containing *K*=1-20 possible sub-populations across 20 replicates, executed in parallel (Mussmann, Douglas, Chafin, et al., 2020). Replicates were clustered and evaluated (Clumpak; Kopelman et al., 2015) to identify potentially discordant modes within a single *K* value. Major clusters were identified using a threshold of 90%. Optimal *K* was chosen by cross-validation (Alexander et al., 2009).

#### 2.3.2 Riverscape genetics using ResistNet

Inputs for ResistNet were derived from the SNP dataset (formatted as VCF) and an input shapefile using autoStreamTree (Chafin et al., 2023). Samples were first grouped by locality and mapped to a stream layer representing all of North America, provided by (RiverATLAS; Linke et al., 2019). Pairwise genetic distances were computed as Weir and Cockerham’s *F*_ST_ (Weir & Cockerham, 1984), linearised (=*F*_ST_/1-*F*_ST_; Rousset; 1997). We subsequently selected a subset of environmental attributes (from N=271; RiverATLAS) as potential covariates for ResistNet. We first excluded those redundant, invariant, or at irrelevant spatial scales for this study, then additionally removed those exhibiting significant collinearity (defined as *r*^2^ > 0.7). This resulted in N=20 environmental and anthropogenic covariates (Table 1).

ResistNet was implemented both globally and individually for each subregion, using 20 independent replicates for each. We used a population size of N=1000, with other settings retained from the simulation-based validation.

## 3. RESULTS

### 3.1 Simulation-based validation

Ensemble models were accurate or contained the correct model in all instances, with exception of sim2 and sim3 for the least topologically complex network (net2; Figure 2), where the correct model was only chosen by 20-60% of replicates (Supplementary Figure S1; Supplementary Table S2). In these cases, ensemble models erroneously excluded SGR (stream gradient; Figure 2). In all instances, convergence occurred in <20 iterations, with a minor sample size effect observed in the delayed convergence for cases more sparsely sampled (i.e., N=10 samples; Supplementary Figure S2). As a composite across simulated scenarios, the correct model was recovered in >90% of replicates. It occurred in100% of ensemble models for net1 when *N*=30 or 50, with lower accuracy at *N*=10 (65% replicates correct and in 75% of ensemble models; Supplementary Table S3).

Accuracies were lowest for net2 (50% of ensemble models correct and 50-75% containing the correct model). For net3, which had the greater spatial extent and topological complexity, 50-75% of ensemble models were correct, but 100% contained the correct model (Supplementary Table S3). Of note, when covariates were erroneously included, they were seemingly selected at a lower weight (Supplementary Figure S3), with much less impact on the model-averaged composite resistances. As above, these cases could generally be identified (and subsequently adjudicated) by examining the consensus, or lack thereof, among replicates and ensemble approaches (Figure 2).

Regarding the methods for summarising replicates, ensemble approaches (ensemble_all and ensemble_best), which employ model-averaging across replicates), offered a means to consolidate discordant replicates, particularly in a few instances where only a minority of replicates recovered the true model (Supplementary Figure S1). As such, they generally performed better when considered across scenarios (Supplementary Table S3).

The most sparsely sampled datasets (*N*=10 samples) had the highest rate of false covariate inclusion (Figure 2), slowest convergence (Supplementary Figure S3), and greatest error in recovering the simulated variable weights (Supplementary Figure S4). They also displayed reduced variation among replicates in model fitness (measured as AIC; Supplementary Figure S5), and often displayed a stronger correlation of the recovered resistance matrix with the input genetic distances (assessed as Mantel’s *r*; Supplementary Figure S6). This suggests a reduced accuracy at smaller sample sizes is not necessarily due to a loss of model optimisation due to insufficient sample, but rather an inability to discriminate amongst multiple model optima which fit equally (or better) than the explicitly simulated conditions. We also note the impact of sample size on covariate selection varied among the three simulated networks, suggesting potential impacts due to spatial dimension, topology, or local conditions relative to the selected covariates.

The most simple of our simulated models [distance-only (sim1) and a two-covariate model with equal weights (sim2)] were most consistently recovered (Figure 2). In contrast, the performance of the models across different network complexities was more nuanced. Here, the least complex topological network (net2) exhibited the lowest accuracy, both in terms of inferred variable importance (Figure 2) and model-averaged weights (Supplementary Figure S3).

### 3.2 *Empirical case study in* Rhinichthys osculus

#### 3.2.1 Assembly and population structure

The global assembly had 13,218 loci with a 25.94x mean depth and 27.6% missing data after filtering. ADMIXTURE identified *K*=30 populations segregating within the five subregions, and broadly corresponding to major tributaries in all cases (Figure 3; Supplementary Figure S6). The Upper Colorado River Basin was divided into *K*=6 groups, while the Virgin River Basin was partitioned into *K*=7. The remaining subregions were divided into *K*=5 linearly distributed through the Grand Canyon, *K*=8 in the Lower Colorado, and *K*=4 in the Little Colorado (Figure 3; Supplementary Figure S6).

**Figure 3:**
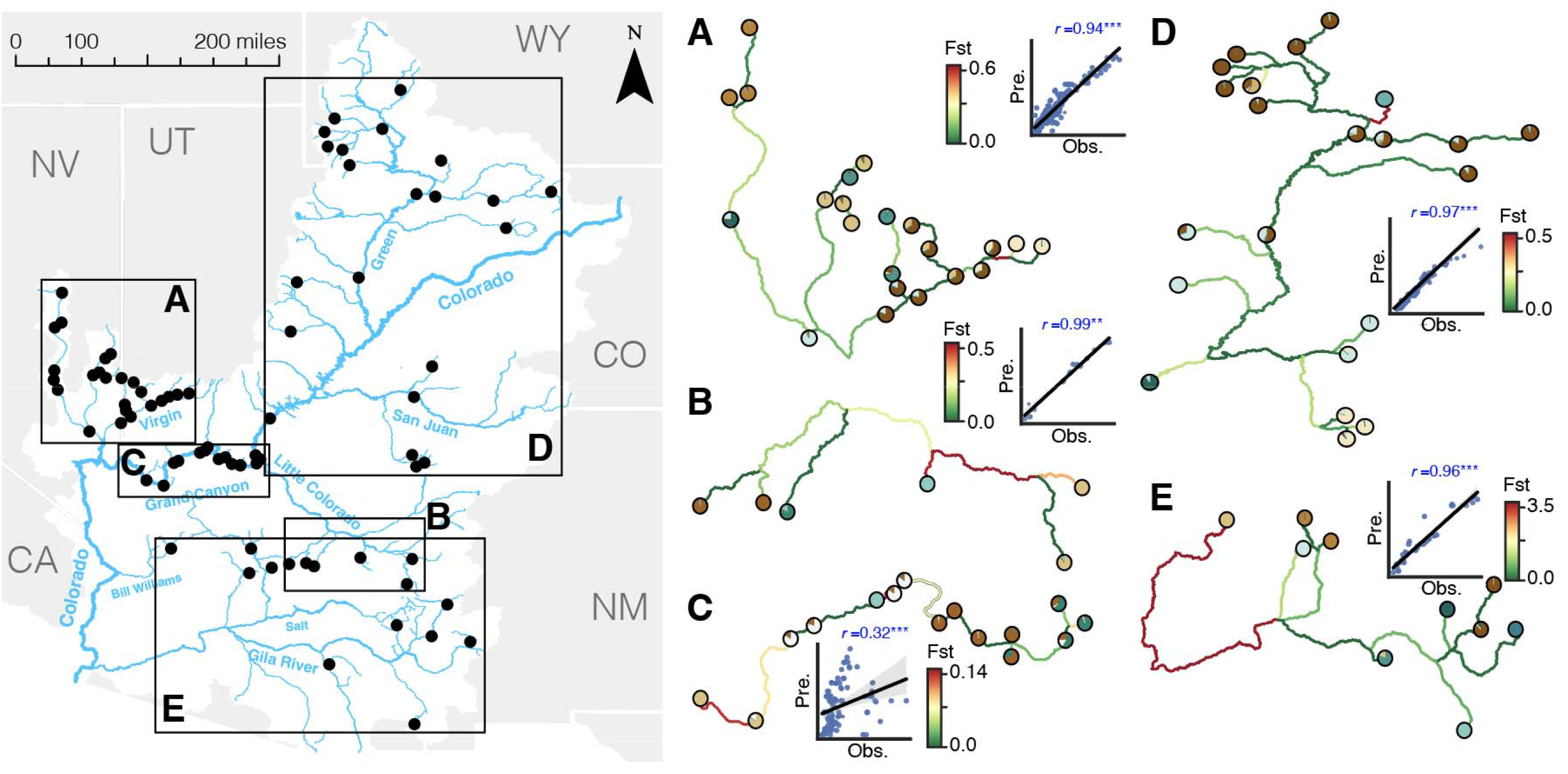
Sampling localities and riverscape genetic structure of *N*=78 sampled populations of Speckled Dace (*Rhinichthys osculus*) in the Colorado River Basin, as divided into five subregions: (A) Virgin River; (B) Little Colorado River; (C) Grand Canyon; (D) Upper Colorado River Basin; (E) Lower Colorado River Basin (Gila and Bill Williams Rivers). Subregion networks are displayed with edges colored by fitted *F*_ST_ (linearised), and sampling locations as mean among-individual assignment probabilities to ancestry clusters derived by ADMIXTURE. Pairwise observed *F*_ST_ (Obs.) is plotted against pairwise fitted *F*_ST_ (Pre.), with significance of Mantel’s *r* indicated at *p*<0.05 (*), *p*<0.01 (**), and *p*<0.001 (***).

#### 3.2.2 Fitted distances and effective resistance networks

The fitted, linearized *F*_ST_ values accurately predicted observed pairwise *F*_ST_ in all subregions (Mantel *r*=0.94-0.99; significant per 1,000 permutations and *p*<0.001), with exception of the Grand Canyon, where correlation was weaker (*r*=0.34) although still significant (*p*<0.001; Figure 3). This indicates that genetic distances mirror network topology, a necessary condition for the appropriate application of ResistNet. As such, ADMIXTURE results generally align well with network topology such that patterns were concordant with fitted, edge-wise *F*_ST_ (Figure 3).

Model-averaged resistances projected onto the stream network (Figure 4) likewise showed a strong concordance with ADMIXTURE probabilities and fitted *F*_ST_ (Figure 3). As such, pairwise resistance significantly predicted *F*_ST_ in all cases (*r*=0.34-0.89) with exception of the Grand Canyon (*r*=0.19). However, resistance models were only marginally better than distance-only models in most subregions, the exception being the Little Colorado (Supplementary Figure S7).

**Figure 4:**
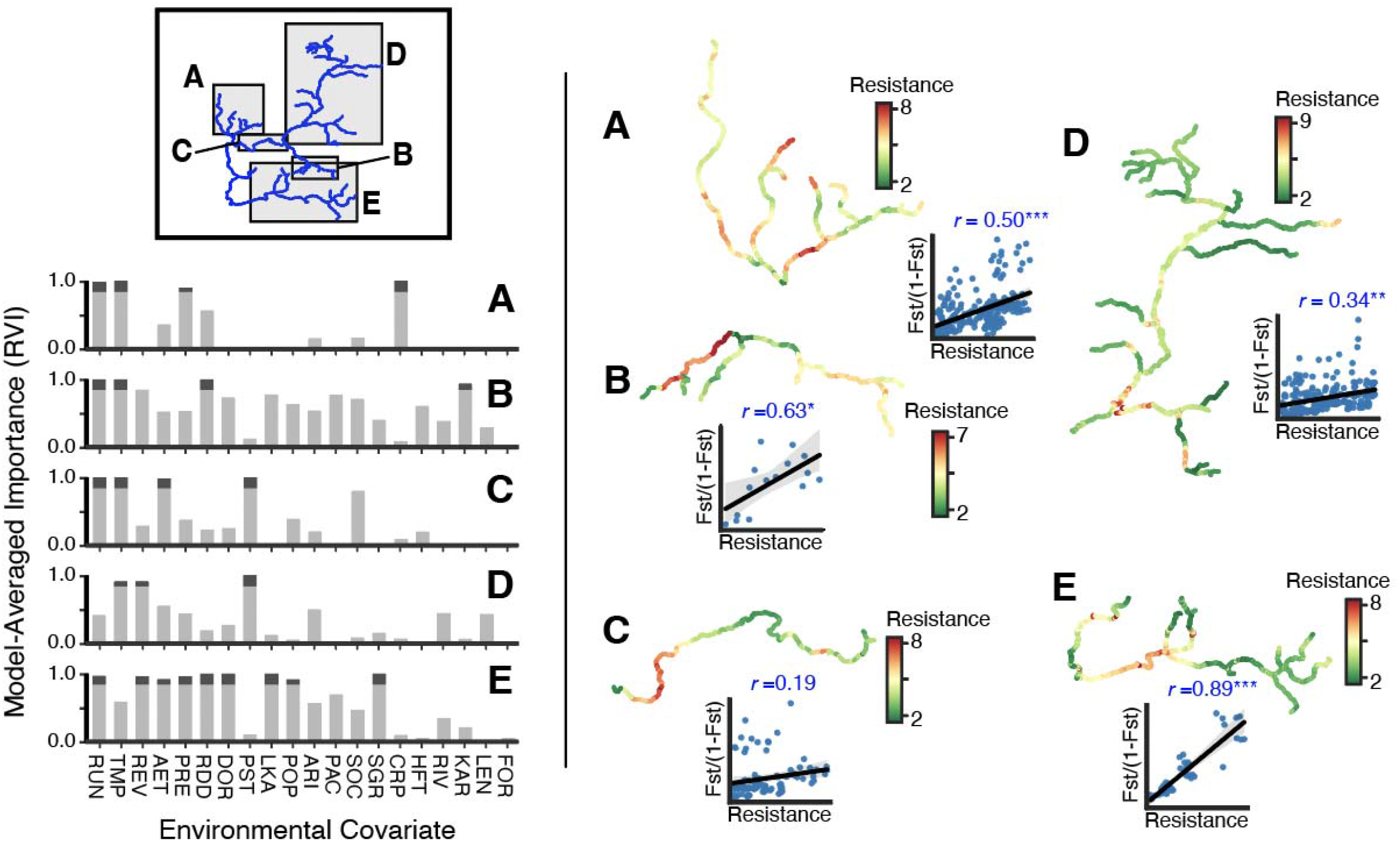
Environmental resistance models and model-averaged resistance networks for Speckled Dace (*Rhinichthys osculus*) from five subregions of the Colorado River. Shown for each subregion (A-E) are model-averaged relative variable importance (RVI) as the sum of Akaike weights for an ensemble model of the best (per AIC) model sampled in each of 20 independent replicates. Here, an RVI threshold of 0.8 is shown in gray. Model-averaged resistance values are plotted on the spatial networks (green=low; red=high), and plotted pairwise against *F*_ST_ [i.e., isolation-by-resistance (IBR)], with significance of Mantel’s *r* indicated at *p*<0.05 (*), *p*<0.01 (**), and *p*<0.001 (***).

Little variation was seen in relative importance (RVI) of covariates when median RVI were compared among replicates using ensemble approaches (Supplementary Figure S8). However, the implicated covariates (using an RVI threshold of 0.8) varied considerably among subregion (Figure 4), with RUN and TMP commonly selected (each chosen in 4/5 subregions).

Specific covariates selected using this threshold for each subregion varied between 3-9: Virgin River (*N*=4; RUN, TMP, PRE, CRP), Little Colorado (*N*=3; RDD, RUN, TMP), Grand Canyon (*N*=4; RUN, TMP, AET, PST), Upper Colorado (*N*=3; TMP, REV, PST), and Lower Colorado (*N*=9; RUN, REV, AET, PRE, RDD, DOR, LKA, POP, SGR). Interestingly, hydrologic distance had minimal contribution across subregions, with a small RVI only present in the Little Colorado and Upper Colorado subregions (Figure 4), but in a minority of replicates therein (Supplementary Figure S8).

The global model, which combined all subregions, selected 14 covariates representing an amalgam of those in the individual subregion models (Supplementary Figure S9). However, effective resistances as model-averages did not show a clear spatial correlation with fitted *F*_ST_ and offered a comparatively weak explanation for observed population assignment probabilities (Figure 3). Given this, we suggest interpretation should be restricted to subregion analyses.

## 4. DISCUSSION

Characterising the spatial distribution of genetic variability within riverscapes serves as a blueprint for the interaction of biodiversity with the environment. Deconstructing this relationship provides a mechanism that can quantify those features which dictate individual movements, thereby informing conservation efforts (Cayuela et al., 2018; Van Strien et al., 2014). To promote this, we introduce ResistNet, a novel method tailored to parse these relationships within spatial networks such as riverscapes. Our simulation-based authentication demonstrates that ResistNet does indeed recover true models of environmental resistance from with high fidelity from pairwise genetic/genomic distances. However, and importantly, our simulated scenarios are idealised, and inherently meet the analytical assumptions underlying the model. Likewise, our case study applied to *Rhinichthys* in the Colorado River Basin underscores both its utility and limitations, which we discuss herein.

As with any method, the applicability of ResistNet must be careful considered within the context of the system, with biological characteristics, the landscape/ riverscape, and the scale and quality of available data being accounted for. Optimization of a resistance model will be poor when ‘true’ correlations among genetic and resistance distances are minimal (Winiarski et al., 2020), or when they are poorly distributed according to network hierarchy (Hopken et al., 2013). This underscores the expectation that topology serves as a reasonable constraint for metapopulation dynamics, a condition unfulfilled for freshwater species with terrestrial or aerial dispersal (Alp et al., 2012; Bohonak & Jenkins, 2003; Phillipsen et al., 2015). Perhaps less obvious are obligate aquatic species that display highly variable dispersal traits and/or behaviours (Comte & Olden, 2018; Tonkin et al., 2018), with emergent patterns of population genetic structure displayed as a cascading effect (Zbinden et al., 2022b). Here, intraspecific genetic differentiation is depicted as a spectrum, even among co-distributed species (Zbinden et al., 2022a), such that it is governed on one hand by differential gene flow within a connected metapopulation, and on the other by isolated patches subject to vicariant processes (Meffe & Vrijenhoek, 1988).

A fundamental assumption of ResistNet, and indeed all methods leveraging the concept of ‘landscape resistance,’ is that individuals act as discrete particles modelled as a ‘random-walk,’ where flux is dictated by resistance values along movement paths that are, by definition, additive, isotropic, and time-invariant (Kumar et al., 2022; B. H. McRae et al., 2008). Emergent patterns of genetic divergence are thus assumed to follow a stepping-stone model (Kimura & Weiss, 1964),that increases as a monotone function of cumulative resistance. However, genetic distances (serving as model input) display a dynamic evolutionary process over non-contemporary timescales. Historic drainage evolution may thus drive cryptic deviations in the model, in that intermittent connections, divergences, and/ or tributary captures may yield genetic patterns at odds with modern hydrography (Douglas et al., 1999). Individual movements via human-mediated translocations will also bias results by altering spatio-genetic associations (Chafin, Zbinden, et al., 2021). It is thus critical to consider both timescale and degree to which the riverscape serves as an evolutionary template (Epps & Keyghobadi, 2015).

Another necessary consideration is the timescale represented by the chosen genetic distance, which must likewise be appropriate for sampling strategy, genetic marker, and the biological context of the study (Landguth et al., 2010; Shirk et al., 2017). Although autoStreamTree supports several alternate distance metrics, such as chord distance (Cavalli-Sforza & Edwards, 1967) or Jost’s *D* (Jost, 2008), we did not evaluate them in a comparative sense as input to ResistNet. It is also important to note that ResistNet implicitly assumes that the contribution of a given environmental feature to effective resistance is spatially homogenous, and that it varies at a scale relevant to the spatial extent and genomic resolution of the study. This may account for the comparatively lower performance of ResistNet in simulations involving our smallest and least topologically complex network (i.e., ‘Net 2’; Figure 2). Smaller sample sizes also carry a greater risk of spurious correlations (Cushman & Landguth, 2010), a trend observable in our simulations as well (Figure 2, Supplementary Table S2, S3). Hence, it is crucial that the sampling design is aligned with both genetic and environmental processes under study (Dauphin et al., 2023). This may translate, for example, to separating samples, thereby more appropriately targeting scale-dependent effects, albeit at a potential loss of statistical power (Keller et al., 2013; Razgour et al., 2014). We explore this topic in the context of our empirical case study below.

Finally, we emphasise the need for environmental data to be properly curated. The scale and accessibility of such data has recently expanded for riverscape (HydroATLAS; Linke et al., 2019)., We recommend this database (and the shapefiles within) as a starting point for rapid exploratory analysis using ResistNet models. We note additional datasets collate probability of intermittence (Messager et al., 2021), and instream barriers (Belletti et al., 2020; Grill et al., 2019) as well. The latter also include several quantitative indices as proxies for the influence of dams (e.g., a factor driving flow regulation). However, the genetic algorithm used by ResistNet can be computationally expensive with a large number of covariates, despite being mitigated through parallel execution. Care should thus be taken to include only explanatory covariates.

Another pervasive confounding factor is multicollinearity (Graham, 2003), which (as above) can vary as a function of sampling configuration (Graves et al., 2013; Prunier et al., 2015). We took a commonly-used pairwise, correlative approach to reduce the number of input features for Rhinichthys (Dormann et al., 2013). Other exclusion techniques taken in this context could involve variation inflation factors (Blair et al., 2013), random forests (Zbinden et al., 2023), or forward-selection based on redundancy analysis (Chafin et al., 2023); all of which could minimise computational complexity by eliminating redundancy in model space, yet with the potential for important data to be reduced (Graham, 2003). Other alternative may seek to either engineer synthetic predictors as combinations of collinear variables, or partition unique *versus* common effects (Dormann et al., 2013; Prunier et al., 2015). Implementation would require pre-processing. Some engineering objectives are integrated within ResistNet as alternative aggregation functions. Although the default is to compute variable-specific resistance as a sum of edge-wise values along the pairwise least-cost path, alternatives such as coefficients of variation or arithmetic/ harmonic means may be selected, though care must be taken that these are appropriate. In summary, effective use of ResistNet requires careful consideration of genetic distance metrics, data treatment, as well as judicious management of trade-offs between variable inclusion and computational expense.

### Empirical case study in Rhinichthys

Our case study not only provides insights specific to our study species, but also general guidelines for applying ResistNet to empirical data such as ours. With regard to drainage-wide population structure, all subregions had reaches with high values of fitted-*F*_ST_ that corresponded with similarly elevated environmental resistance (Figure 3, 4), to include those representing relatively well-connected riverscapes. IBR was generally a better predictor than (IBD (Figure 4, Supplementary Figure S7). Hydrologic distances were likewise non-explanatory as a component of resistance models, implicating environmentally-mediated connectivity as more important for intraspecific divergences. This, in turn, complements analyses of population structure (Figure 3, Supplementary Figure S6), and juxtaposes well with prior observations designating unique and relictual lineages (Mussmann, Douglas, Oakey, et al., 2020; Oakey et al., 2004).

Numerous covariates strongly implicated for *Rhinichthys* were linked to connectivity though flow, drought tolerance, and seasonal persistence of streams (e.g., RUN, TMP, AET, PRE), suggests a role for flow regime/ ephemerality (Figure 4). Unsurprisingly, hydrology is a driver of morphological diversification in numerous freshwater systems (Bruckerhoff & Magoulick, 2017; Langerhans, 2008; Meyers & Belk, 2014), and its importance in sustaining native fish assemblages has been documented across southwestern streams (Kominoski et al., 2018; Propst et al., 2008; Propst & Gido, 2004). The large weights attributed to variables that define stream persistence clearly reflect the impacts of aridity and persistent drought, both of which are omnipresent across contemporary desert riverscapes (Comte et al., 2022).

Covariates relating to contemporary (e.g., anthropogenic) effects such as impoundment and flow regulation were also chosen (Figure 4). Here, factors relating to anthropogenic land use [e.g., road density (RDD); pasture extent (PST); population density (POP)]; lake/ reservoir extent [limnicity (LKA); upstream reservoir volume (REV)], and indices of hydrologic manipulation [degree of regulation (DOR)], most strongly contributed to composite resistance models. Several are functionally related, such as: Impoundment as a mechanism for flow regulation/ flood dampening (DOR), and determining reservoir volume (REV), channel width, and morphology (Grams et al., 2020; Walker et al., 2020). Anthropogenic factors promoting habitat instability in the Colorado River have previously been implicated as modulating species boundaries in other endemic leuciscids (Chafin et al., 2019). These altered conditions may also change the distribution and suitability of low flow refugia, serving in turn to extirpate peripheral populations (Bezzerides & Bestgen, 2002; Magoulick & Kobza, 2003; Schlosser, 1995). Altered conditions also promote the establishment of non-native species (Dibble et al., 2021; Gido & Propst, 1999; Whitney et al., 2017), which may in turn elevate predation rates and promote competition (Marsh & Douglas, 1997; Seegert et al., 2014). Here, predation pressure has been shown to curtail invasion success of *Rhinichthys osculus* post-introduction (Harvey et al., 2004), thus suggesting a potential role for non-natives in biotic resistance.

An observation of broader relevance to the applicability of ResistNet for *Rhinichthys* is the relative fit of optimised resistance networks and, by extension, the applicability of IBR varies substantially as an explanatory model by scale and subregion. The basin-wide analysis, in particular, was a poor fit (Supplementary Figure S9), which we hypothesized as reflecting the transection of historical or vicariant processes not captured by extant connectivity. This is evidenced by the apparent ‘plateau’ in the global relationship of *F*_ST_ with hydrologic distance (Supplementary Figure S9b), suggesting the relative importance of genetic drift and gene flow as being scale dependent (Hutchison & Templeton, 1999; Keller et al., 2013). Here, inflated genetic divergences may also emerge as vestiges of historic barriers, such as Pleistocene lava flows within the Grand Canyon region (Duffield et al., 2006).

Broad-scale hydrologic evolution, as driven by tectonism and vulcanism, are a hallmark of the region (Minckley et al., 1986), with genetic vestiges also well-characterised in other co-occuring species (Chafin, Douglas, et al., 2021). More recent population dynamics could also lead to a situation where genetic divergences fail to juxtapose with extant conditions, a possibly explanation of high divergences displayed by Vermillion Creek, which may have inadvertently escaped large-scale, indiscriminate rotenone poisoning as a mechanism to exterminate non-native species in the Green River in the 1950s-60s (Holden, 1991). Finally, it is also possible that environmental and/ or anthropogenic components of riverscape resistance may vary at smaller spatial scales, especially in species widely distributed.

Another confounding factor could be admixture involving unsampled or distant sites (Jaquiéry et al., 2011). For example, intergradation between morphological forms of *Rhinichthys osculus* in the Little Colorado and Lower Colorado subregions has been hypothesised as being driven by historic stream capture along the Mogollon Rim, with historically-transient connections possible as well between some Upper Colorado River Basin tributaries and the Bear or Snake rivers (Loxterman & Keeley, 2012; Smith & Dowling, 2008). Historic reticulation violates an assumption of our resistance model, in that connectivity does not involve edges represented by the network. If such effects are indeed unrecognized, then *F*_ST_-values fitted along that stretch would be incorrectly inflated. Our strategy herein was to define units for analysis from apparent population structure (Mussmann, 2018), phylogeny (Oakey et al., 2004), and a biogeographic understanding of the region (Minckley et al., 1986). However, quantitative approaches to construct input matrices could employ an iterative resampling procedure (Goldberg & Waits, 2010) to identify appropriate spatial thresholds or topological constraints (Savary et al., 2021).

### Conclusion

IBR may not be appropriate for all systems – and indeed, our *Rhinichthys* case study demonstrates the potential for intraspecific variation – and explanatory models may thus be conflated or wholly redundant (Zbinden et al., 2022a). However, ResistNet can achieve very high accuracies (>95%), when scale is appropriately considered along both spatial and topological dimensions. These limitations serve as a necessary context from which to develop additional applications, or serve as avenues for future research, as do emerging developments in riverscape genetics (White et al., 2020). One extension for locus-wise analysis (e.g., Chafin et al., 2023), is to identify functional genomic associations that can promote isolation-by-adaptation rather than environmental resistance or limitation via dispersal (Orsini et al., 2013). Adaptive genetic variation is a relatively underexplored theme in riverscape genetics (Davis et al., 2018), due largely to a dearth of methods for detecting ‘outlier’ loci against the backdrop of a unique autocorrelative structure of networks.

Another advantage of ResistNet is the potential to extrapolate and predict genetic responses in unsampled space. Here, optimized resistance models may be applied across any dendritic ecological network (or riverscapes, as herein), to predict effective resistance in regions within which genetic data has yet to be obtained. This could promote *a priori* hypotheses of population and phylogeographic structure that can supplement transecting regions with elevated divergences or inform analyses requiring assignment priors (Hubisz et al., 2009). It could also extrapolate connectivity as an invasive (Lovell et al., 2021), or project temporal changes due to predicted climate change or other disturbance (Nakajima et al., 2023). Importantly, the approach can be applied to any type of pairwise distance matrix. For example, autoStreamTree could be applied to examine edge-wise distances or topological importance in generating differences within community data (Eros et al., 2011), with ResistNet then applied to examine how environmental resistance defines metacommunity structure. Again, any form of spatially structured ecological network can be generalized for such accommodation.

The potential of ResistNet to accelerate automation in riverscape genetics can also encourage large-scale comparative riverscape studies involve variation among species in spatial or temporal trends, the results of which could vary substantially as a function of trait synergism or life histories (Comte et al., 2014; McManamay & Frimpong, 2015; Troia et al., 2019). While rare, such comparative studies are critical for testing ecological and evolutionary questions (Blanchet et al., 2020; Fourtune et al., 2016; Paz-Vinas et al., 2018; Pilger et al., 2017; Zbinden et al., 2022b, 2023). Further, a comparative framework also allows those commonalities driving community-level structure or shared biogeographic breaks to clarify, which may supplement conservation planning at contemporary spatial or taxonomic scales (Linke et al., 2011; Paz-Vinas et al., 2018), and effectively promote multi-species management objectives (Nielsen et al., 2017).

## ACKNOWLEDGEMENTS

Computational resources were provided by the Arkansas High Performance Computing Center (AHPCC). MED and MRD are supported by the University of Arkansas via endowments (MED: 21^st^ Chair Century Global Change Biology; MRD: Bruker Life Sciences Professorship). TKC is supported by the Scottish Government’s Rural and Environment Science and Analytical Services Division (RESAS). We also thank the members of numerous state, federal, and tribal entities throughout the Colorado River Basin who provided tissue samples which enabled this research. The use of trade, product, industry, or firm names is for informative purposes only and does not constitute an endorsement by the U.S. Government or the U.S. Fish and Wildlife Service. Links to non-Service websites do not imply any official U.S. Fish and Wildlife Service endorsement of the opinions or ideas expressed therein or guarantee the validity of the information provided. The findings, conclusions, and opinions expressed in this article represent those of the authors, and do not necessarily represent the views of the U.S. Fish & Wildlife Service.

## AUTHOR CONTRIBUTIONS

All authors contributed to study design; SMM, MRD, and MED performed lab work and collected data; SMM performed bioinformatics and assembly; TKC wrote software and designed methodology; SMM and TKC analysed the data; TKC drafted the initial manuscript. All authors contributed critically to the drafts, interpretation of results, and approved of the manuscript in its final form.

## DATA AVAILABILITY

All source code for ResistNet is available on GitHub: https://github.com/tkchafin/resistnet. Simulated and empirical datasets used to validate ResistNet, including Snakemake workflows to replicate analyses, are available via the Open Science Framework: https://osf.io/4uxfj. Raw sequence reads are deposited in the NCBI SRA under BioProject PRJA656098.

---

**Figure S1:**
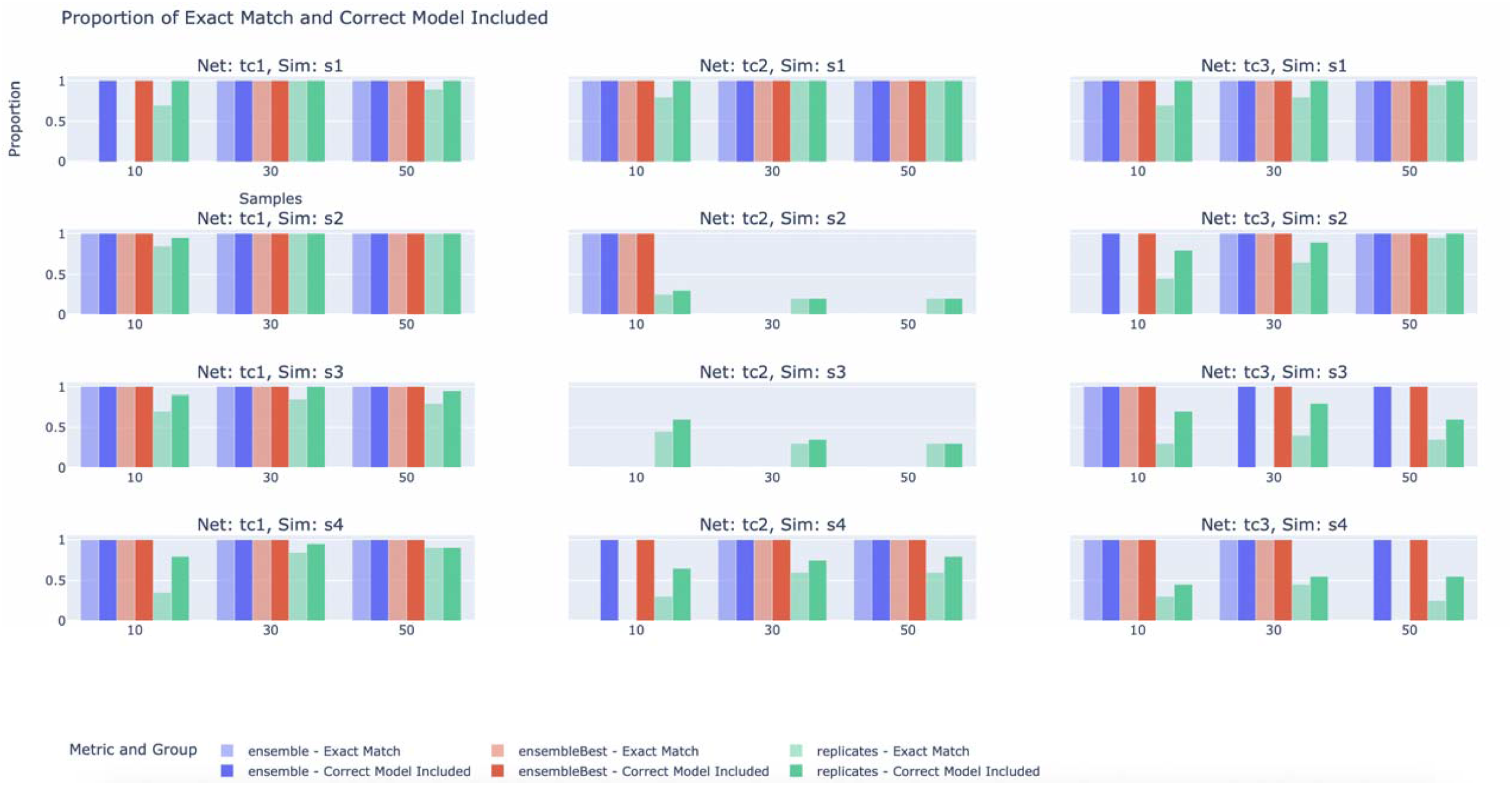
Validation of ResistNet using three networks and four simulated scenarios. The proportion of replicates (green) is shown, illustrating the model-averaged result that accurately recovered the known conditions (light green) in contrast to those containing the known condition along with additional spurious covariates (dark green). Results are also presented for across-replicate ensemble models (blue) and ensemble models constructed exclusively from the best AIC models selected for each replicate (referred to as “ensembleBest”; red). Note that the results are further divided by sample size (10, 30, or 50).

**Figure S2:**
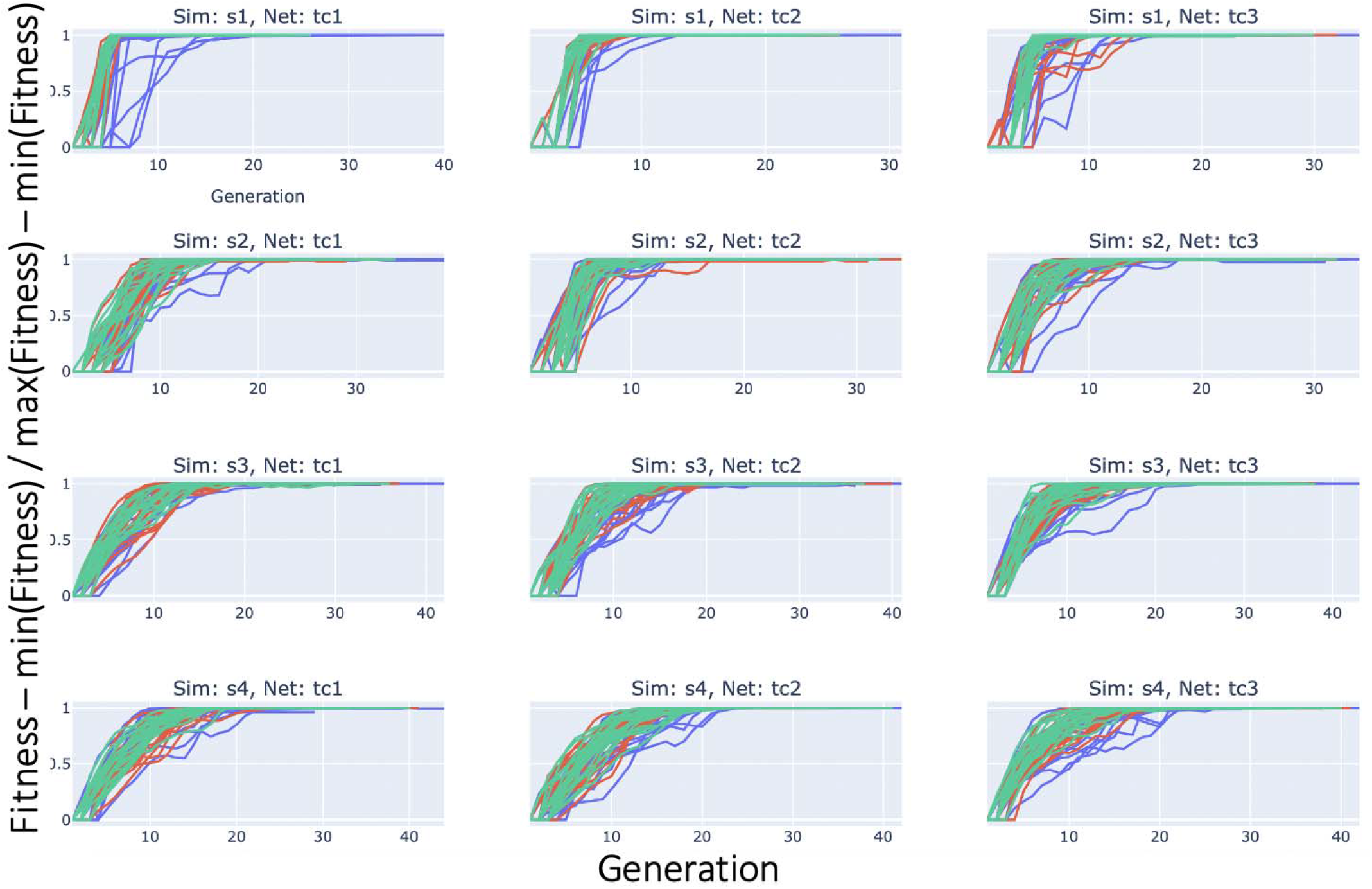
Model optimization results for simulation-based validation of ResistNet conducted on three networks and four simulated scenarios. Results are presented for simulations with 10 (blue), 30 (red), and 50 (green) samples, each performed in 20 independent replicates. The fitness values (Y-axis) represent AIC, shown as absolute values transformed onto a consistent scale across replicates using min-max normalisation.

**Figure S3:**
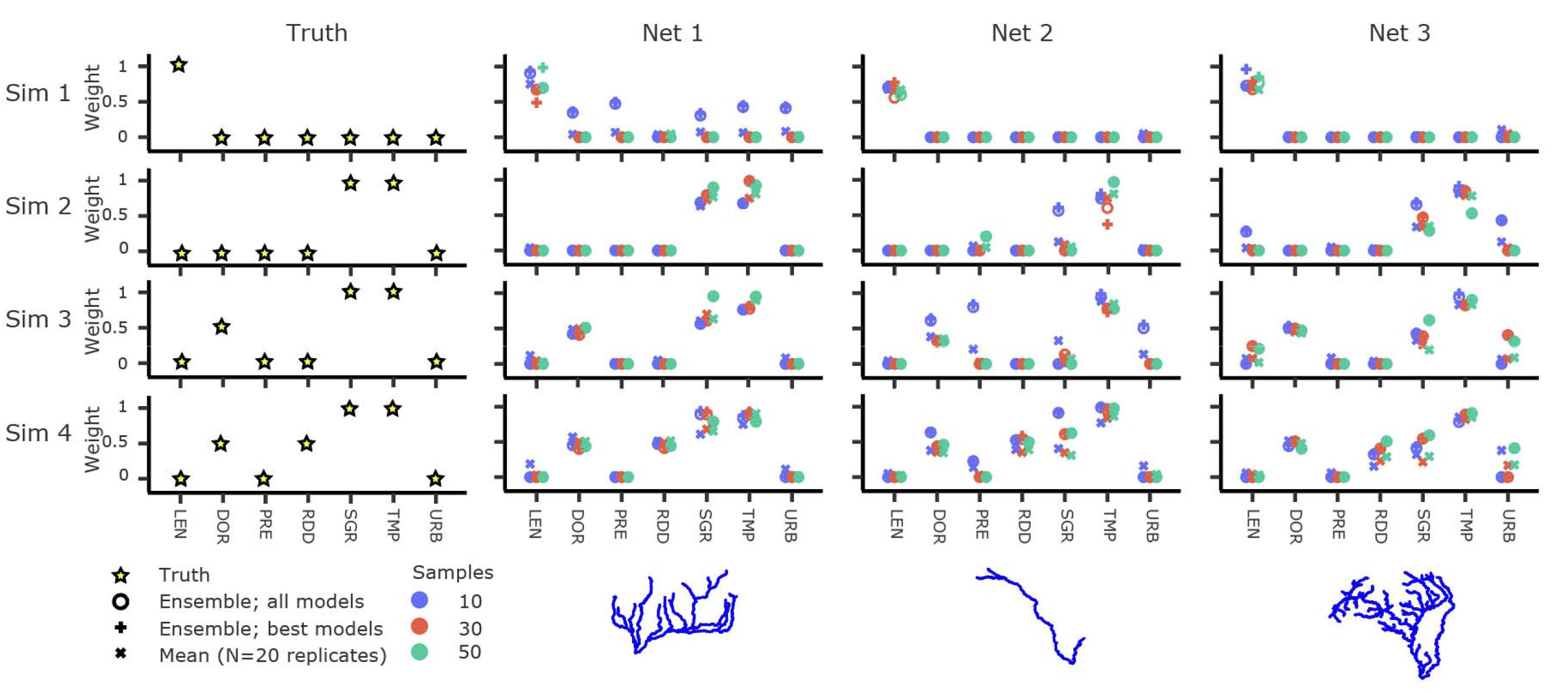
Summary of model-averaged weights (MAW) from simulated scenarios. MAWs were derived from four pre-determined ‘known’ scenarios (Sim1-4), tested within three distinct template networks (Net1-3). Results are reported as: (1) Ensemble (all): a MAW computed from all models chosen across 20 independent replicates; (2) Ensemble (best): MAW calculated exclusively from the ’optimum’ models in each replicate, determined by the Akaike Information Criterion (AIC); and (3) Median: The mean MAW amongst all replicates. Each scenario was modeled using three different sample sizes: 10, 30, and 50, represented by blue, red, and green points, respectively.

**Figure S4:**
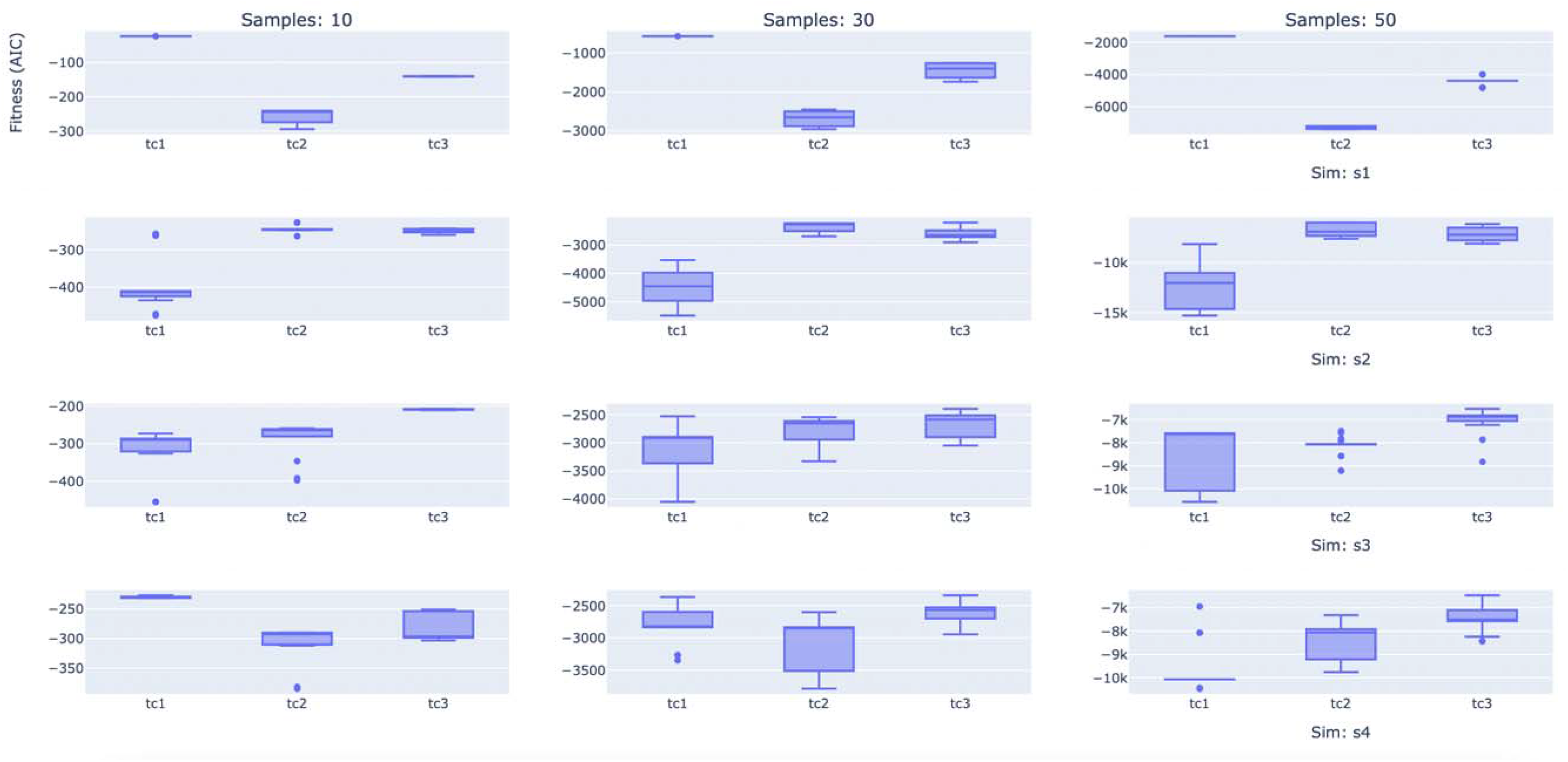
AIC values across replicates for simulation-based validation of ResistNet, performed on three networks and four simulate scenarios. Results are partitioned by simulation scenario (rows) and sample size (columns; 10, 30, or 50 samples), with results shown by network (x-axis).

**Figure S5:**
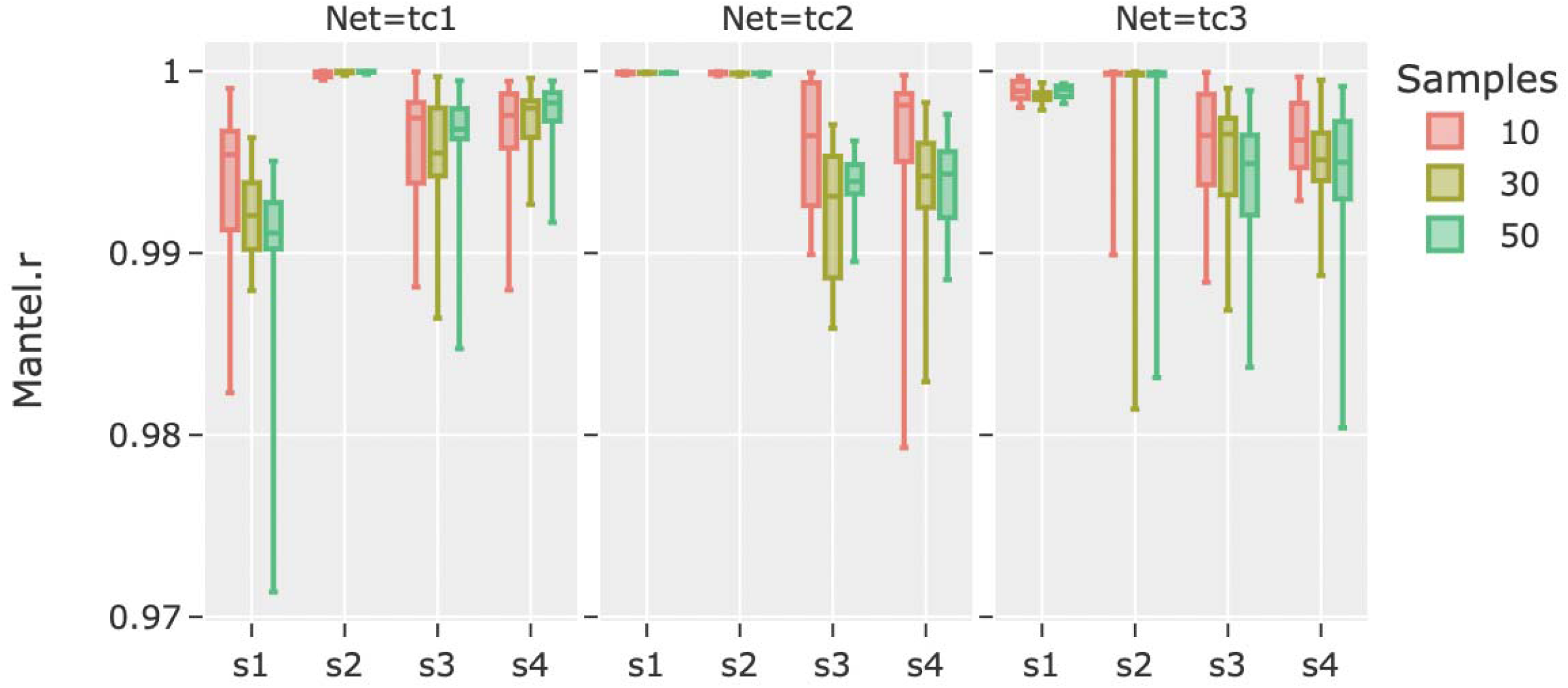
Correlation of output pairwise resistance matrices among replicate ResistNet runs (*N*=20) with simulated input distance matrices across four simulated scenarios (s1-4) run within three template networks (tc1-3), at varying sample sizes (10, 30, 50), reported as Mantel’s *r*.

**Figure S6:**
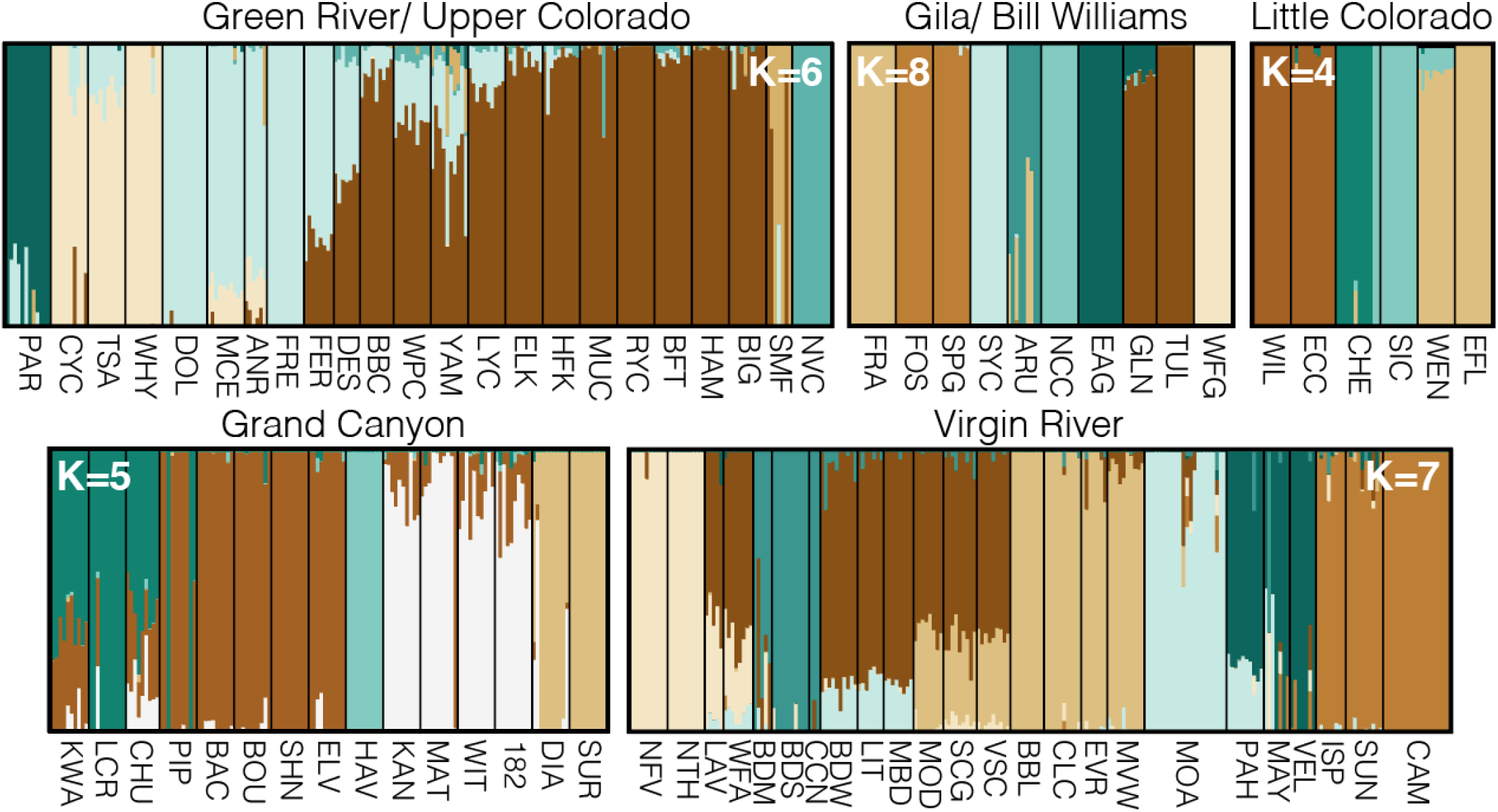
Population structure among Colorado River Basin (CRB) Speckled Dace populations. Upper Colorado/ Green River aggregates (*K*=6) represent: Paria River (PAR), Chinle Wash (CYC-WHY), San Juan+San Rafael+Fremont rivers (DOL-FER), Lower-Mid Green River (DES-BIG), Smiths Fork (SMF), and Vermillion Creek (NVC). The Lower CRB, represented by the Gila+Bill Williams River drainages (*K=*8) include: the Bill Williams River (FRA), Verde River (FOS-SPG), Agua Fria (SYC), San Pedro (ARU), San Simon (NCC), Eagle Creek (EAG), San Francisco River (GLN-TUL), and Upper Gila River (WFG). The Little Colorado River (*K*=4) encompasses East Clear Creek (WIL-ECC), Chevelon Creek (CHE), Silver Creek (SIC), and the East Fork (WEN-EFL). Grand Canyon (*K*=5): distributed linearly along the length of the canyon, with the exception of Havasu Creek (HAV). Virgin River (*K*=7): upper Virgin River (NTH-NFV), Mainstem Virgin River (LAV-WFA; BDW-VSC), Beaver Dam Wash (BDM-CCN), Meadow Valley Wash (BBL-MVW), Moapa River (MOA), Pahranagat (PAH-VEL), and White River (ISP-CAM).

**Figure S7:**
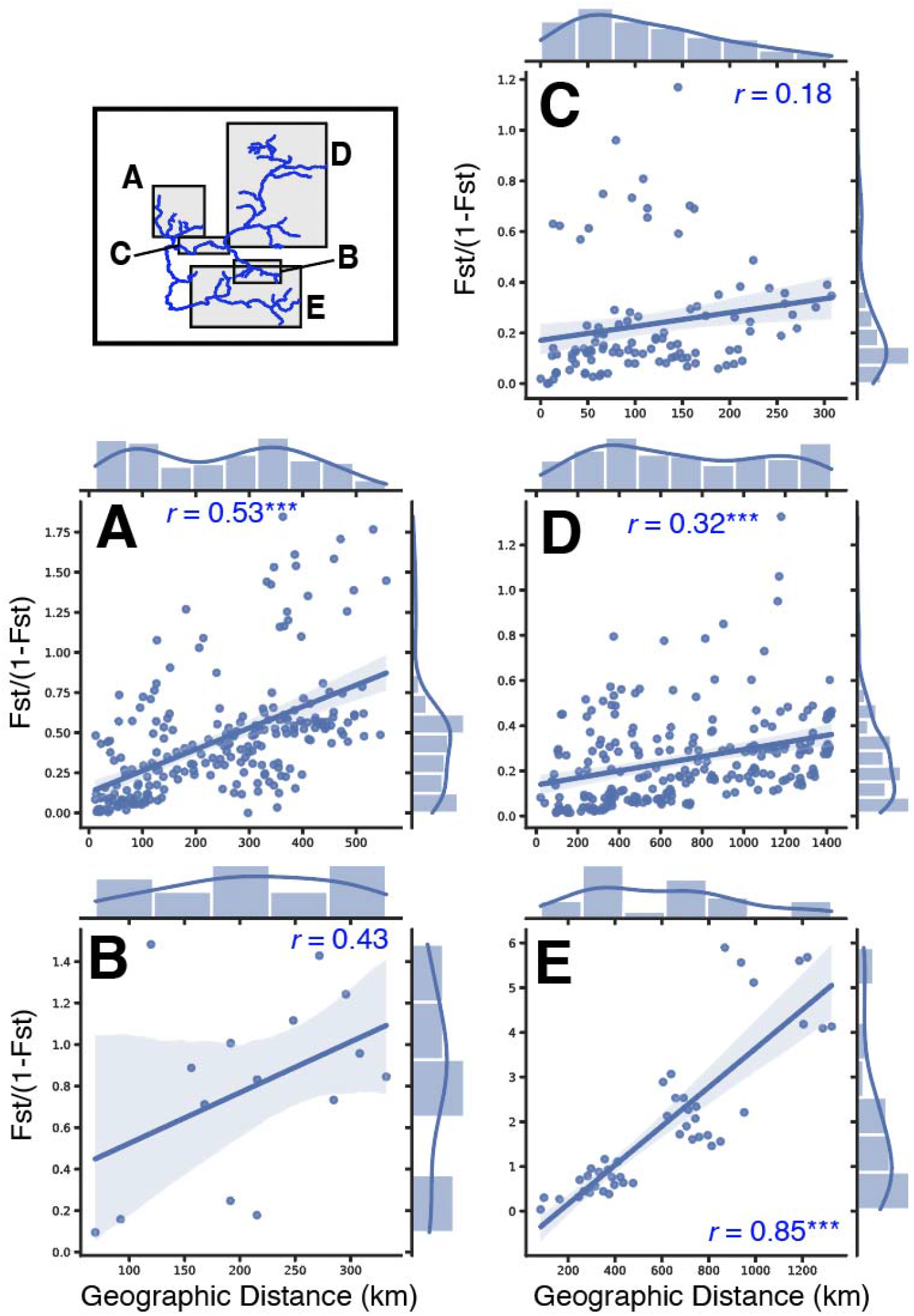
Isolation-by-distance (IBD) for Speckled Dace (*Rhinichthys osculus*) populations sampled from five subregions of the Colorado River Basin, plotted as the pairwise geographic/ hydrologic distance (in km) and linearised *F*_ST_. Correlation is reported as Mantel’s *r*, with significance indicated at *p*<0.05 (*), *p*<0.01 (**), and *p*<0.001 (***).

**Figure S8:**
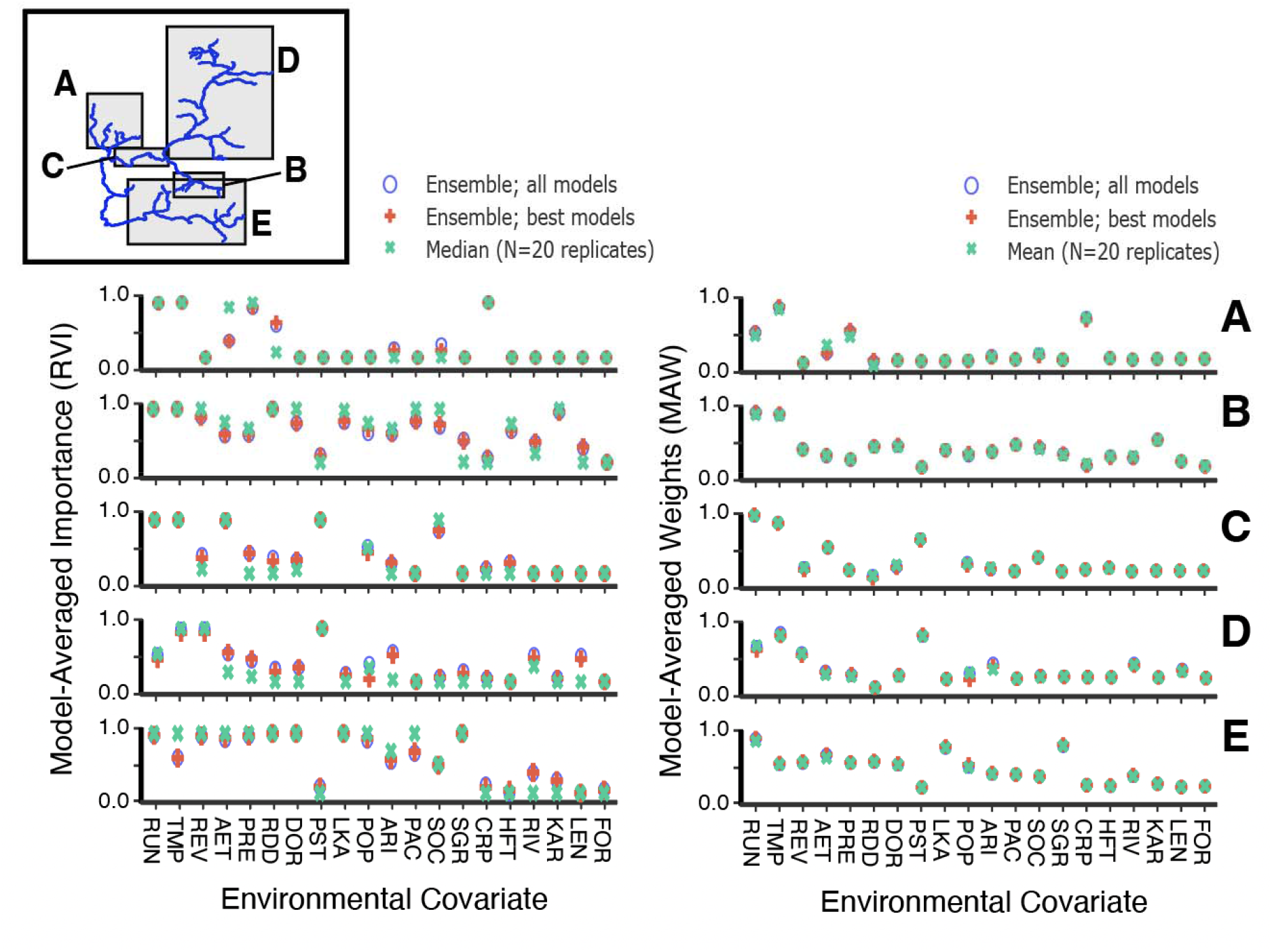
Model-averaged relative variable importance (RVI) and weights (MAW) for environmental resistance models derived using ResistNet applied to Speckled Dace (*Rhinichthys osculus*) from five subregions of the Colorado River. Shown for each subregion (A-E) are RVIs and MAWs reported from an ensemble of all sampled models (ensemble_all; blue), an ensemble of the best model sampled within each replicate (ensemble_best; red), and as the mean (MAW) or median (RVI) of replicates (green).

**Figure S9:**
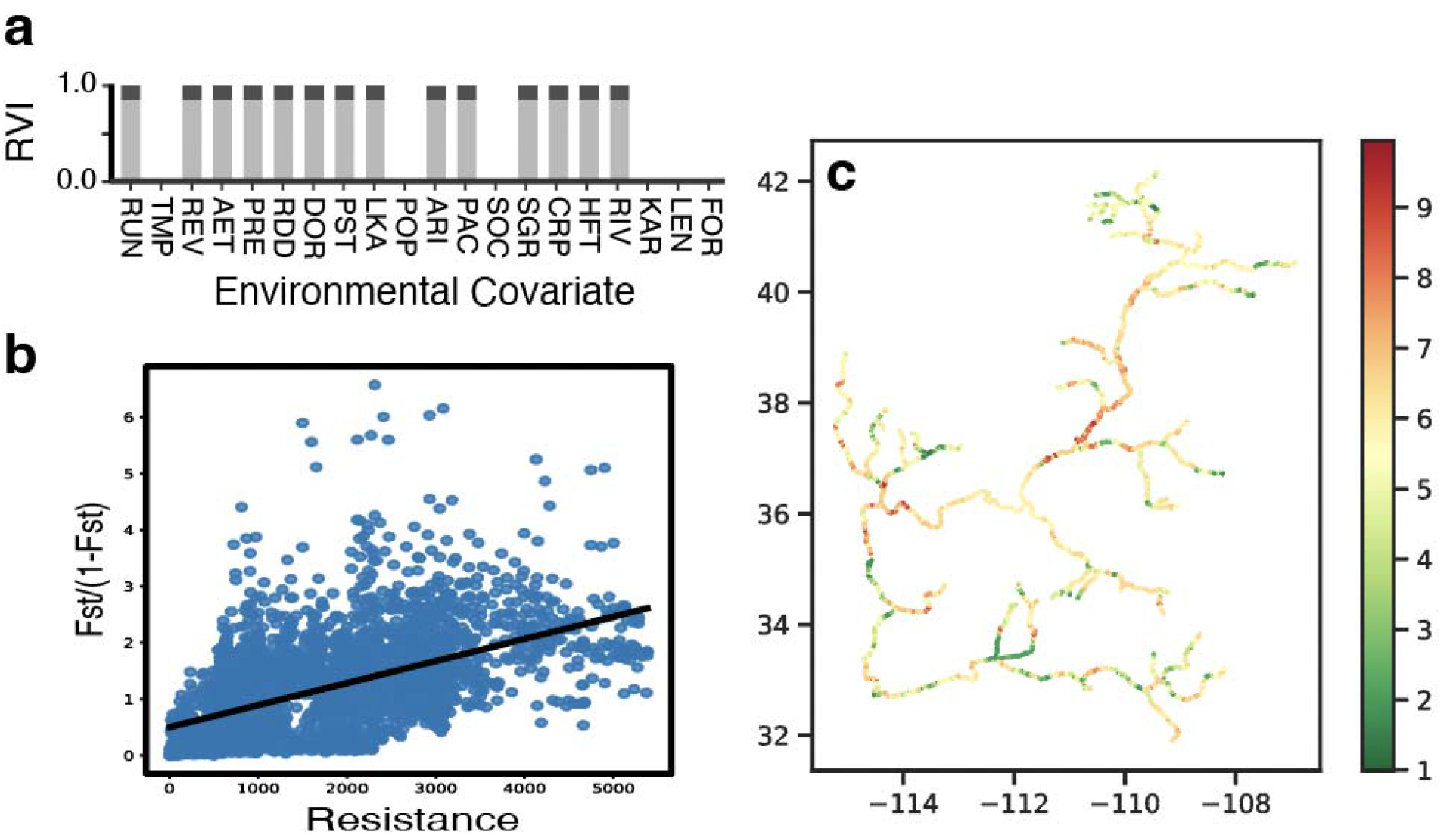
Environmental resistance models and model-averaged resistance network for Speckled Dace (*Rhinichthys osculus*) evaluated globally across *N*=78 sampling localities in the Colorado River. Shown are model-averaged relative variable importances (RVIs) as the sum of Akaike weights for an ensemble model of the best (per AIC) model sampled in each of 20 independent replicates. Here, an RVI threshold of 0.8 is shown in gray. Model-averaged resistance values are plotted on the spatial networks (green=low; red=high), and plotted pairwise against *F*_ST_ [i.e., isolation-by-resistance (IBR)].

**Table S1:** Locality information (=Locality) and sample sizes (=*N*) for *N*=78 sampling localities of Speckled Dace (*Rhinichthys osculus*) in subregions of the Colorado River Basin.

| Site | Locality | Subregion | N |
| --- | --- | --- | --- |
| 182 | Grand Canyon RM182 | Grand Canyon | 10 |
| ANR | Animas River | Upper CRB | 6 |
| ARU | Aravaipa Creek; San Pedro tributary | Lower CRB | 12 |
| BAC | Grand Canyon RM87-117 | Grand Canyon | 10 |
| BBC | Big Beaver Creek – White River | Upper CRB | 9 |
| BBL | Meadow Valley Wash | Virgin | 9 |
| BDM | Beaver Dam Wash | Virgin | 6 |
| BDS | Beaver Dam Wash | Virgin | 10 |
| BDW | Mainstem Virgin River | Virgin | 10 |
| BFT | Blacks Fork | Upper CRB | 10 |
| BIG | Big Sandy River | Upper CRB | 10 |
| BOU | Grand Canyon RM87-117 | Grand Canyon | 10 |
| CAM | Camp Spring – White River | Virgin | 18 |
| CCN | Beaver Dam Wash | Virgin | 4 |
| CHE | Chevelon Creek; Little Colorado trib. | Little Colorado | 12 |
| CHU | Grand Canyon RM225-249 | Grand Canyon | 9 |
| CLC | Meadow Valley Wash | Virgin | 10 |
| CYC | Chinle Wash; Colorado R. tributary | Upper CRB | 10 |
| DES | Desolation Canyon; Green R. mainstem | Upper CRB | 7 |
| DIA | Grand Canyon RM225-249 | Grand Canyon | 10 |
| DOL | Dolores River | Upper CRB | 12 |
| EAG | Eagle Creek | Lower CRB | 12 |
| ECC | East Clear Creek; Little Colorado trib | Little Colorado | 12 |
| EFL | East Fork, Little Colorado River | Little Colorado | 10 |
| ELK | Elk River (Yampa River Tributary) | Upper CRB | 10 |
| ELV | Grand Canyon RM87-117 | Grand Canyon | 10 |
| EVR | Meadow Valley Wash | Virgin | 7 |
| FER | Ferron Creek – San Rafael River | Upper CRB | 8 |
| FOS | Fossil Creek; Verde River tributary | Lower CRB | 10 |
| FRA | Francis Creek; Bill Williams R. tributary | Lower CRB | 12 |
| FRE | Fremont River; Green River tributary | Upper CRB | 10 |
| GLN | San Francisco River at Glenwood | Lower CRB | 9 |
| HAM | Ham's Fork; Green River | Upper CRB | 10 |
| HAV | Havas Creek, Grand Canyon | Grand Canyon | 10 |
| HFK | Ham's Fork; Green River | Upper CRB | 10 |
| ISP | Indian Springs; White River; Virgin-Muddy tributary | Virgin | 8 |
| KAN | Kanab Creek; Grand Canyon | Grand Canyon | 10 |
| KWA | Grand Canyon RM225-249 | Grand Canyon | 10 |
| LAV | Mainstem Virgin River | Virgin | 5 |
| LCR | Little Colorado River confluence | Grand Canyon | 10 |
| LIT | Mainstem Virgin River | Virgin | 7 |
| LYC | Little Yampa Canyon; Yampa River | Upper CRB | 10 |
| MAT | Grand Canyon RM143-189 | Grand Canyon | 10 |
| MAY | Maynard Spring – Pahrnagat Valley | Virgin | 7 |
| MBD | Mainstem Virgin River | Virgin | 8 |
| MCE | McElmo Creek – San Juan River | Upper CRB | 10 |
| MOA | Moapa River; Virgin-Muddy tributary | Virgin | 22 |
| MOD | Mainstem Virgin River | Virgin | 8 |
| MVW | Meadow Valley Wash | Virgin | 10 |
| MUC | Muddy Creek – Blacks Fork River | Upper CRB | 10 |
| NCC | San Simon | Lower CRB | 10 |
| NFV | Upper Virgin River | Virgin | 10 |
| NTH | Upper Virgin River | Virgin | 10 |
| NVC | Vermillion Creek | Upper CRB | 10 |
| PAH | Pahrnagat River; Virgin-Muddy basin | Virgin | 10 |
| PAR | Paria River | Upper CRB | 12 |
| PIP | Grand Canyon RM87-117 | Grand Canyon | 10 |
| RYC | Ryckman Creek upstream of Hill Creek Confluence | Upper CRB | 10 |
| SCG | Mainstem Virgin River | Virgin | 9 |
| SIC | Silver Creek; Little Colorado tributary | Little Colorado | 10 |
| SHN | Shinumo Creek; Colorado R. tributary | Grand Canyon | 10 |
| SMF | Smith’s Fork; Green River | Upper CRB | 7 |
| SPG | Spring Creek; Verde River tributary | Lower CRB | 10 |
| SUN | Sunnyside Creek; White River; Virgin-Muddy | Virgin | 10 |
| SUR | Grand Canyon RM225-249 | Grand Canyon | 10 |
| SYC | Sycamore Creek; Agua Fria tributary | Lower CRB | 10 |
| TSA | Chinle Wash; Colorado R. tributary | Upper CRB | 10 |
| TUL | Tularosa Creek – San Francisco River | Lower CRB | 10 |
| VEL | Burn’s Ranch – Pahrnagat Valley | Virgin | 7 |
| VSC | Mainstem Virgin River | Virgin | 10 |
| WEN | East Fork, Little Colorado R. | Little Colorado | 10 |
| WFA | Mainstem Virgin River | Virgin | 8 |
| WFG | West Fork of the Gila River at cliff dwellings | Lower CRB | 10 |
| WIL | East Clear Creek; Little Colorado trib | Little Colorado | 10 |
| WIT | Grand Canyon RM143-189 | Grand Canyon | 10 |
| WPC | Island Park below Whirlpool Canyon; Yampa River | Upper CRB | 10 |
| WHY | Chinle Wash; Colorado R. tributary | Upper CRB | 10 |
| YAM | Yampa River (Green R. tributary) | Upper CRB | 10 |

**Table S2:**
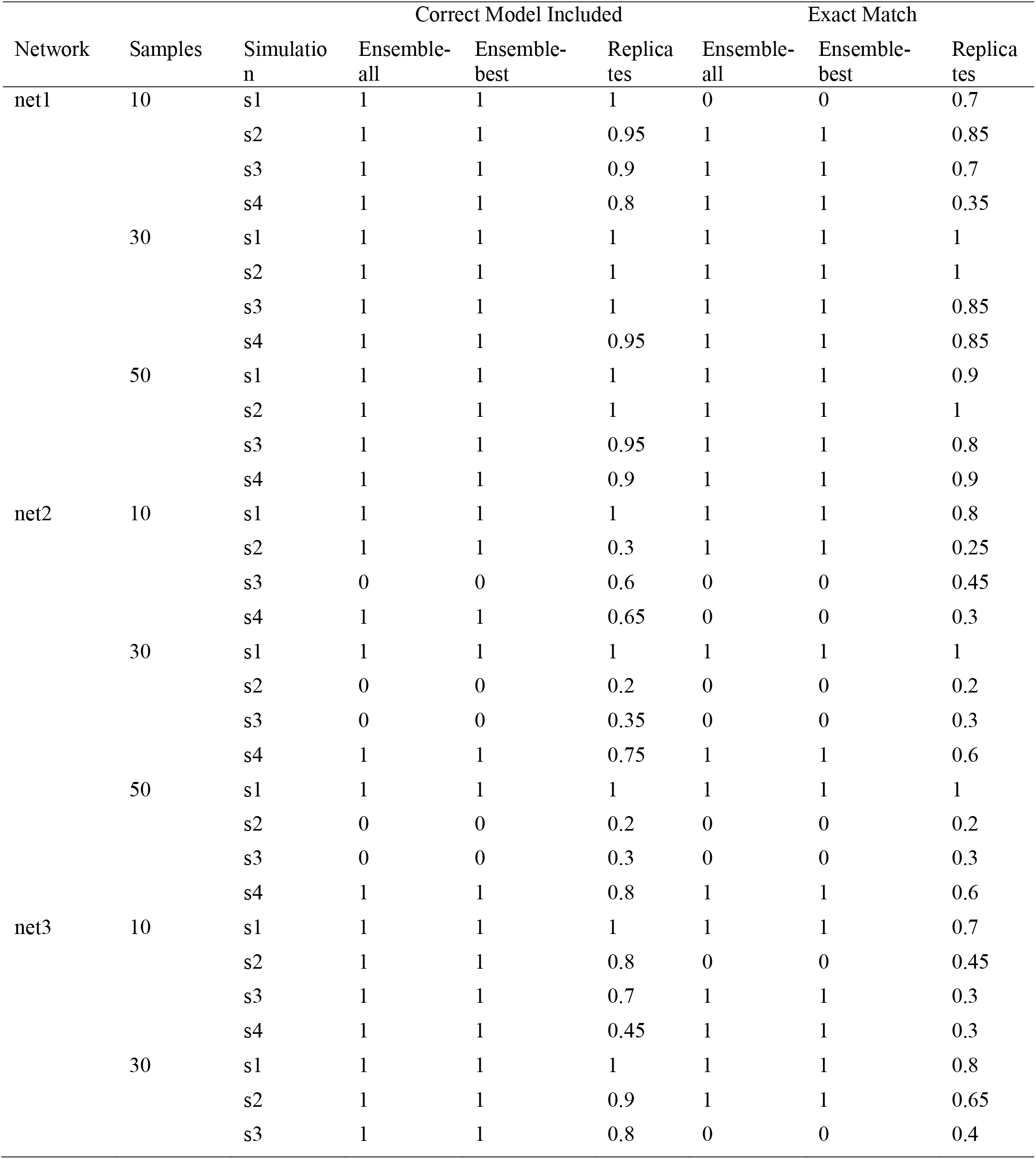
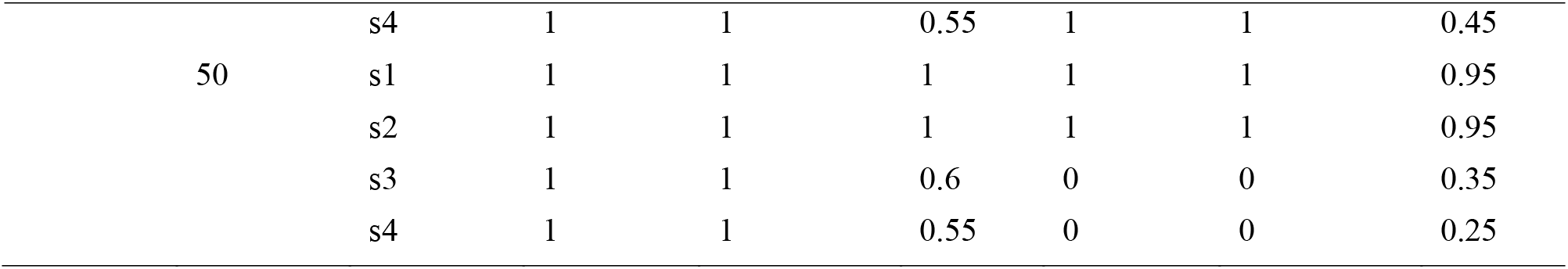
Accuracy of ResistNet validations runs performed using four simulated scenarios (s1-s4) on three template networks (net1-3), and three sample sizes (10, 30, 50). Shown for each are the proportion of replicates for which the model-averaged output accurately captured the exact simulated scenario (“Exact Match”), or encapsulated the simulated scenario (e.g., having an additional covariate implicated; “Correct Model Included”). The same metrics are also shown for the ensemble_all and ensemble_best approaches, but as true (1) or false (0).

**Table S3:**
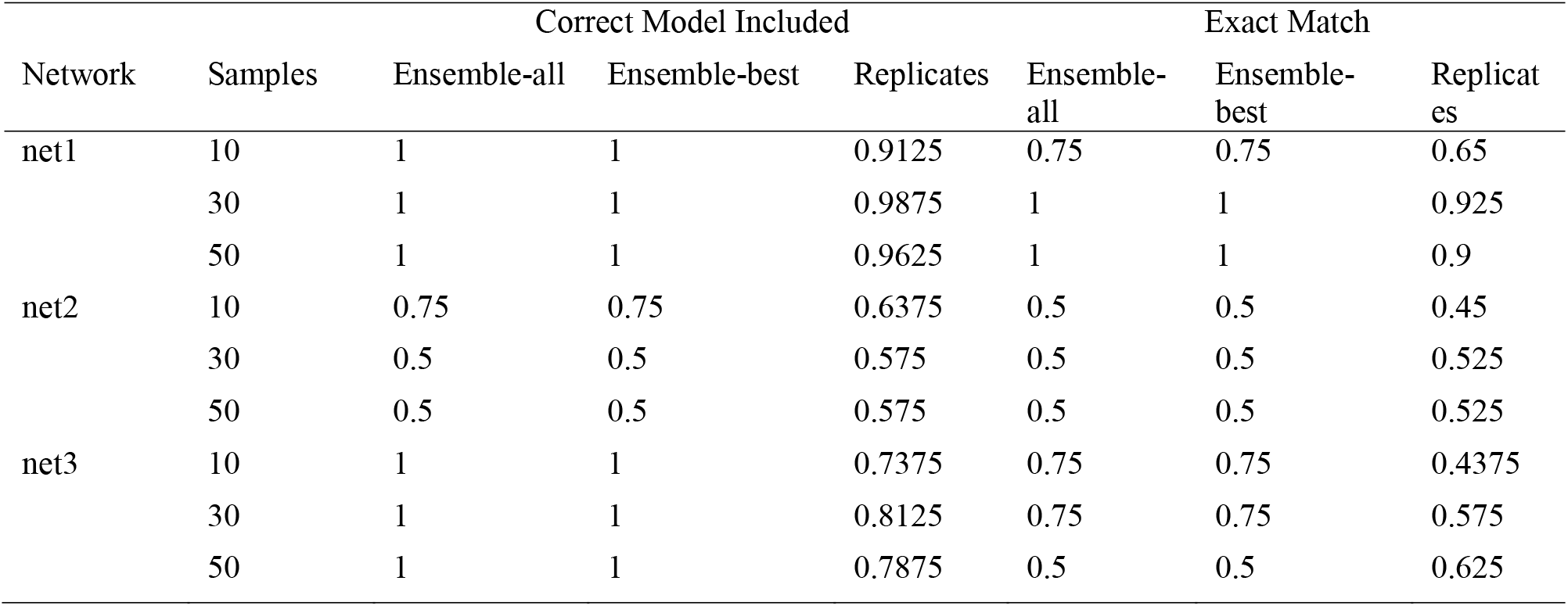
Accuracy of ResistNet validations runs performed using four simulated scenarios (s1-s4) on three template networks (net1-3), and three sample sizes (10, 30, 50). Shows are results summarised across all simulated scenarios, with the proportion of replicates or ensemble models (ensemble_best and ensemble_all) for which the model-averaged output accurately captured the exact simulated scenario (“Exact Match”), or encapsulated the simulated scenario (e.g., having an additional covariate implicated; “Correct Model Included”).

